# Sub-millisecond conformational dynamics of the A_2A_ adenosine receptor revealed by single-molecule FRET

**DOI:** 10.1101/2020.11.26.400184

**Authors:** Ivan Maslov, Oleksandr Volkov, Polina Khorn, Philipp Orekhov, Anastasiia Gusach, Pavel Kuzmichev, Andrey Gerasimov, Aleksandra Luginina, Quinten Coucke, Andrey Bogorodskiy, Valentin Gordeliy, Simon Wanninger, Anders Barth, Alexey Mishin, Johan Hofkens, Vadim Cherezov, Thomas Gensch, Jelle Hendrix, Valentin Borshchevskiy

## Abstract

The complex pharmacology of G-protein-coupled receptors (GPCRs) is defined by their multi-state conformational dynamics. Single-molecule Förster Resonance Energy Transfer (smFRET) is well-suited to quantify dynamics for individual protein molecules, however, its application to GPCRs is challenging; therefore, smFRET has been limited to studies of interreceptor interactions in cellular membranes and receptors in detergent environments. Here, we performed smFRET experiments on functionally active human A_2A_ adenosine receptor (A_2A_AR) molecules embedded in freely diffusing lipid nanodiscs to study their intramolecular conformational dynamics. We propose a dynamic model of A_2A_AR activation that involves a slow (>2 ms) exchange between the active-like and inactive-like conformations in both apo and antagonist-bound A_2A_AR, explaining the receptor’s constitutive activity. For the agonist-bound A_2A_AR, we detected faster (390±80 μs) ligand efficacy-dependent dynamics. This work establishes a general smFRET platform for GPCR investigations that can potentially be used for drug screening and/or mechanism-of-action studies.

## Introduction

G-protein-coupled receptors (GPCRs) constitute the largest superfamily of membrane proteins in humans containing over 800 members, which mediate critical physiological processes, such as neurotransmission, homeostasis, inflammation, reproduction, olfaction, vision, taste, and others^1,2^. GPCRs recognize a large variety of endogenous extracellular signaling molecules transmitting their corresponding signals inside the cell, and this process can be modulated by synthetic ligands or drug molecules. In fact, over 30% of all FDA-approved drugs target GPCRs^3^. Multiple lines of evidence suggest that the molecular mechanism of GPCR activation extends beyond a simple ‘on/off mode. First, apo receptors show basal activity that can be suppressed by inverse agonists^4^. Second, different agonists vary in efficacy and can stimulate receptor activity to a different extent^5^. Third, a single receptor can signal through several intracellular pathways, some of which could be preferentially activated by so-called “biased” ligands^6^. These three phenomena indicate that receptors are highly dynamic molecules and sample several active and inactive states stochastically (for review, see^7–9^).

The A_2A_ adenosine receptor (A_2A_AR) is expressed in many organs and tissues including those in the immune system, basal ganglia, heart, lungs, and blood vessels^10^. Throughout the body, A_2A_AR regulates the cardiovascular tonus causing vasodilation and promotes healing of inflammation-induced injuries by suppressing immune cells^11,12^. In the brain, A_2A_AR modulates dopamine and glutamate neurotransmission^12^. A_2A_AR is a promising target for drugs against insomnia, chronic pain, depression, Parkinson’s disease, and cancer^12,13^. On the molecular level, A_2A_AR is activated by the endogenous extracellular agonist adenosine and initiates the cAMP-dependent signaling pathway via G_s_ and G_olf_ proteins^12,14^. Besides G-proteins, A_2A_AR interacts with numerous other partners including GRK-2 kinase, β-arrestin, and other GPCRs^14,15^. One cryoEM and over 50 high-resolution X-ray crystallographic structures are available for antagonist- or agonist-bound A_2A_AR and for its ternary complex with an agonist and an engineered G protein, making this receptor an excellent model system for investigating GPCR structural dynamics. While static structures provide critical information about the receptor’s lowest energy states, our understanding of the A_2A_AR function remains critically incomplete without the detailed knowledge of its conformational dynamics.

The current information about A_2A_AR conformational dynamics is based mostly on several reported NMR experiments^16–22^. In response to ligand binding, different A_2A_AR amino acids either alter their sole stable conformations or vary relative probabilities of coexisting stable conformations^16,17^. On the picosecond-to-nanosecond timescale, some A_2A_AR amino acids increase side-chain dynamics, while others become stabilized^18^. Sub-millisecond conformational variability was shown for both apo-form^19^ and agonist-bound A_2A_AR^16,17,20^. Large-scale conformational changes in A_2A_AR with dwell times of seconds were also reported^19,21^, but two independent studies described the corresponding long-lived states differently: in one report^19^, a 3-state model with an attributed basal activity of 70% was proposed, while in the other, the authors put forward a 4-state model with a negligible basal activity. Thus, the current picture of A_2A_AR dynamics is complex and contradictory.

Studies of A_2A_AR dynamics face two major challenges: first, the need to cover a wide range of timescales from nanoseconds to seconds, and next, the difficulty to untangle multiple protein states within the ensemble. Single-molecule fluorescence spectroscopy provides tools to address both of these difficulties. Depending on the applied method, the fluorescence signal from individual receptors can be tracked with as low as a nanosecond temporal resolution for a total duration of either milliseconds in case of freely-diffusing molecules or even seconds to minutes using immobilized molecules^7,23^.

Single-molecule fluorescence spectroscopy methods have been previously applied to studying GPCR conformational dynamics^7^. For example, environmentally-sensitive fluorescent dyes have been used as single-molecule reporters of conformational changes in the β_2_ adrenergic receptor (β_2_AR)^24–27^ and visual rhodopsin^28,29^. Single-label experiments are attractive because of a minimal influence of the dye on the native receptor dynamics, but the experimental readouts are often limited and lack detailed structural interpretation. Additionally, the results of single-label experiments can be obscured by multi-state dye photophysics. Another approach, based on single-molecule Förster Resonance Energy Transfer (smFRET) between two dyes can provide more direct structural outcomes and introduce additional internal controls, however, at the expense of double-labeling. smFRET has been shown to be especially useful to investigate structural dynamics of GPCR dimers^30–34^. To our knowledge, at the moment of this writing, ref.^35^ is the only published application of smFRET to quantifying intramolecular conformational dynamics in GPCRs; this study addressed structural changes on the intracellular side of immobilized β_2_AR in detergent micelles.

Here we applied smFRET to investigate the conformational dynamics of A_2A_AR in lipid nanodiscs freely diffusing in solution without immobilization. Using the MFD-PIE (multiparameter fluorescence detection with pulsed interleaved excitation) technique^36^ (Fig. 1A), we tracked the relative movements of two dyes attached to the intracellular tip of the transmembrane helix TM6 (L225C^6.27^, superscripts indicate Ballesteros–Weinstein numbering^37^) and to the C-terminal intracellular helix H8 (Q310C^8.65^) of A_2A_AR (Fig. 1B). We observed that FRET efficiency in the double-labeled A_2A_AR increases upon agonist-binding (Fig. 1C). Several burst-wise fluorescence analysis approaches — plot of burst-wise FRET efficiency against donor fluorescence lifetime^38^, FRET 2-Channel kernel-based Density Estimator (FRET-2CDE)^39^, Burst Variance Analysis (BVA)^40^, and filtered Fluorescence Correlation Spectroscopy (fFCS)^41^ – subsequently revealed sub-millisecond conformational dynamics of A_2A_AR. Based on quantitative analysis of the obtained data for the receptor in its apo state and upon addition of the inverse agonist ZM241385, the partial agonist LUF5834, or the full agonist NECA to the receptor, we finally propose a dynamic model of A_2A_AR activation.

**Fig. 1.**
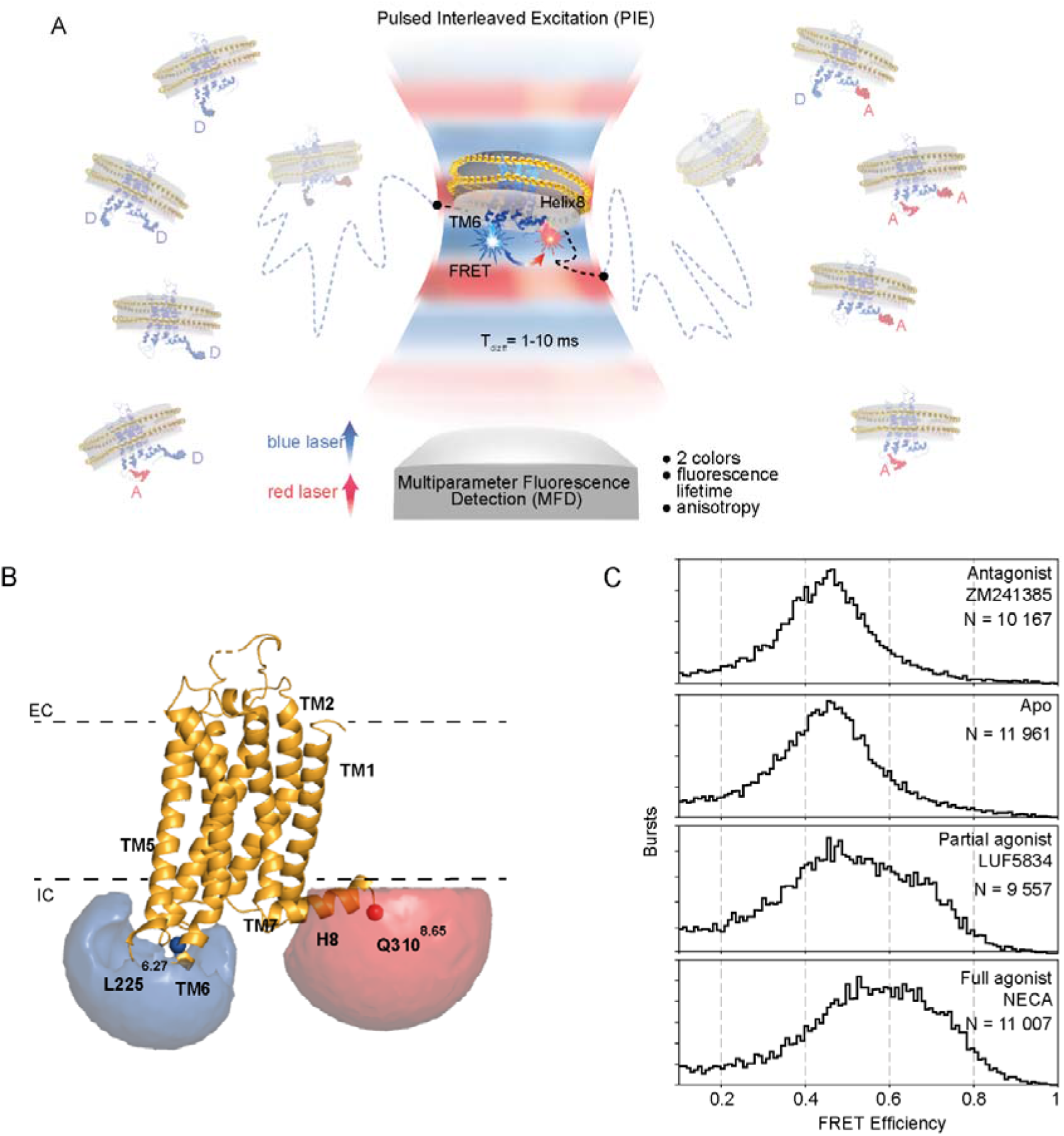
Agonist-induced conformational changes in A_2A_AR are revealed by smFRET. **(A)** Schematic illustration of the MFD-PIE smFRET experiment on A_2A_AR embedded in lipid nanodiscs and stochastically labeled with the donor (Alexa488) and the acceptor (Atto643) fluorescent dyes at TM6 and H8. Eight coexisting labeling variants of A_2A_AR are shown as shadowed receptors in both sides of the image, ‘D’ and ‘A’ correspond to donor and acceptor dyes, respectively. A_2A_ARs diffuse in solution and stochastically cross the focal spot of an inverted fluorescence microscope. Bursts of fluorescence from donor and acceptor fluorophores are recorded within the 1-10 ms residence time of individual A_2A_ARs crossing the focal spot. Only those receptors labeled with both donor and acceptor produce FRET signal. In the PIE approach, two spatially overlapped and alternatingly pulsing lasers are focused by the microscope objective to excite donor and acceptor fluorescence consequently. Using the MFD approach, fluorescence signals of donor and acceptor are recorded separately, and the fluorescence lifetime and anisotropy of each dye are determined. **(B)** The labeled sites (L225^6.27^, Q310^8.65^) and the volume accessible for the dyes (simulated using FPS software^78^) are shown on the A_2A_AR structure (PDB: 3EML^58^), the membrane boundaries (dashed lines) are obtained from the PPM web server^79^ and shown as dashed lines **(C)** Burst-wise distributions show an agonist-induced increase in FRET efficiency in the double-labeled A_2A_AR. The number of bursts used for the analysis (N) is given for each condition.

## Results

### Labeling and reconstitution of A_2A_AR in nanodiscs

To track the conformational dynamics of A_2A_AR with smFRET we chose to attach two fluorescent dyes to mutated residues L225C^6.27^ on the intracellular end of TM6 and Q310C^8.65^ on the C-terminal end of H8 (Fig. 1B, Fig. S1A). In previous A_2A_AR FRET studies, a fluorescent protein-based FRET donor and fluorescent molecule based acceptor in similar labeling positions were shown to provide sufficient contrast between the active and inactive receptor states in live cells^42,43^. The residue position L225^6.27^ is also homologous to the native cysteine C265^6.27^ in β_2_AR that has been frequently used for fluorescent labeling^24–26,44–48^.

We expressed the double-Cys mutant (L225C^6.27^/Q310C^8.65^) of A_2A_AR in *Leishmania tarentolae* and simultaneously labeled it with two maleimide-functionalized dyes, Alexa488 and Atto643. The wild-type (WT) A_2A_AR has six unpaired cysteines in its transmembrane helices (Fig. S1A). To achieve specific labeling of the two genetically introduced cysteines, but spare the transmembrane native cysteines, we labeled the receptors in isolated cell membranes, as described previously^49^. After labeling, the receptors were purified and reconstituted in MSP1D1 nanodiscs, which can accommodate only a single monomeric receptor per nanodisc^50^.

Size-exclusion chromatography confirmed a high purity and monodispersity of the nanodisc-reconstituted A_2A_AR samples (Fig. S1B). Labeling efficiencies of 26% (Alexa488) and 8% (Atto643) were obtained for the mutant A_2A_AR (Table S1). Labeling specificity was confirmed with the WT receptor, which showed only a marginal dye fluorescence associated with the protein after the labeling procedure (Fig. S1B). In both ensemble spectra and lifetime measurements of the fluorescently labeled A_2A_AR FRET-sensitized acceptor emission was readily observed, proving the existence of double-labeled FRET-active molecules in the samples (Fig. S1C-D).

To test whether the double-cysteine mutant A_2A_AR (L225C^6.27^/Q310C^8.65^) is functional, we measured the ligand-induced thermostabilization of the isolated receptors as well as the agonist-induced cAMP accumulation in living cells. A fluorescent thermal stability assay^51^ showed that the addition of either the antagonist ZM241385 or the agonist NECA in saturating concentrations increased the melting temperature of both WT and mutant A_2A_AR with respect to the apo-state by >7 ° C, indicating ligand-binding activity of the receptor (Fig. S1E). A BRET assay of cAMP accumulation in HEK293T cells transiently expressing A_2A_AR showed very similar pEC_50_ values for both WT (6.41±0.15) and double-mutant (6.45±0.06) forms of the receptor upon stimulation with the agonist NECA (Fig. S1F). The mutant form of A_2A_AR retained functional activity in nanodiscs and in cells, therefore we assume that the conformational dynamics observed for the double-labeled receptor in smFRET experiments represent the native dynamics of the WT receptor.

### smFRET reveals ligand-induced conformational changes in A_2A_AR

We diluted fluorescently labeled A_2A_AR to single-molecule concentrations, mounted the sample on a microscope cover slip and recorded fluorescence intensity, lifetime, and anisotropy data from individual molecules diffusing freely across the femtoliter-sized observation spot (approximated by a 3D Gaussian with half-widths 0.5 μm, 0.5 μm and 2 μm) of a confocal fluorescence microscope (Fig. 1A). Inside the spot, donor and acceptor fluorophores are excited alternatingly using a two-color pulsed-interleaved excitation (PIE)^52^. The residence time of individual molecules (~1-10 ms) in the laser spot sets the upper limit of timescales approachable for the observation of A_2A_AR conformational dynamics. Using a 4-detector MFD scheme (Fig. S2), photons detected from individual molecules were digitally tagged with (1) the spectral band in which they were detected, (2) their global arrival time with microsecond accuracy, (3) their relative arrival time with respect to the laser pulses within a ps-ns range, and (4) their optical polarisation^53^. PIE together with two-color detection allowed us to distinguish double-labeled receptors (simultaneously labeled with donor and acceptor) from “donor-only” and “acceptor-only” receptors (Fig. S3, see ‘Selection of double-labeled, donor-only and acceptor-only subpopulations’ in SI).

The fraction of A_2A_ARs simultaneously labeled with donor and acceptor fluorophores showed different distributions of FRET efficiency depending on the bound ligand (Fig. 1C). The antagonist ZM241385 did not change FRET efficiency distribution within experimental error. On the contrary, both the partial agonist LUF5834 and the full agonist NECA shifted the mean FRET efficiency to larger values and increased the overall distribution width, compared to the apo-receptor. The increase in FRET efficiency was less pronounced for the partial agonist LUF5834 than for the full agonist NECA.

### Fluorescence lifetime data suggest sub-millisecond conformational dynamics of A_2A_AR

Besides fluorescence intensity, FRET is also reflected in fluorescence lifetime data. A two-dimensional plot of the per-burst FRET efficiency against the donor fluorescence lifetime provided the first insights into the receptor’s conformational dynamics (Fig. 2A,B). In theory, data for rigid molecules, in which FRET efficiency remains constant over the duration of a burst should be distributed along a curved diagonal line that intersects the lifetime axis at the lifetime of the donor-only population and the FRET efficiency axis at unity, commonly referred to as the ‘static FRET line’ (Fig. 2A). Alternatively, if receptor molecules sample different conformations during their residence time in the focal spot (1-10 ms) on a timescale that is longer than the nanosecond fluorescence lifetime, their bursts should be shifted from the ‘static FRET line’ towards the longer lifetime region. This phenomenon can be explained by the higher weights of the lower FRET states in the fluorescence lifetime averaging due to the larger number of photons emitted by the donor. The observed rightward deviations of our burst data from the static FRET line indicate the existence of sub-millisecond conformational dynamics (beyond the fast dynamics expected for dye linkers) in the apo as well as agonist- and antagonist-bound states of A_2A_AR (Fig. 2B).

**Fig. 2.**
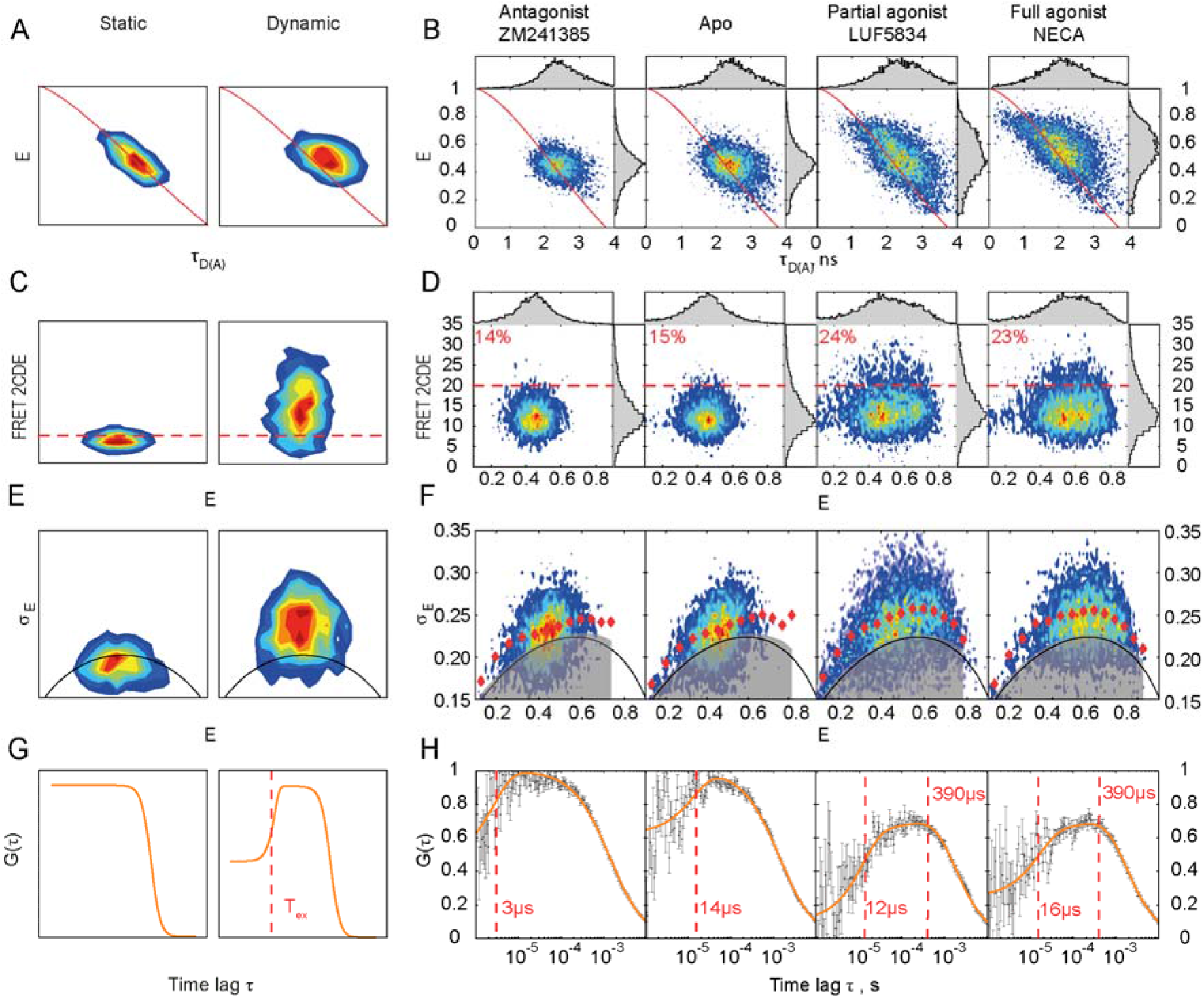
Four complementary burst-wise analysis approaches suggest an agonist-induced increase in the sub-millisecond conformational dynamics of A_2A_AR. Contour plots are two-dimensional histograms of different fluorescence burst parameter distributions. Simulated plots for ‘static’ and ‘dynamic’ molecules **(A,C,E,G)**, and experimental data for double-labeled A_2A_AR (**B,D,F,H**) are shown. **(A, B)** The FRET efficiency is plotted against donor fluorescence lifetime. The ‘static FRET’ line is shown in red. A shift of burst disctibution to the right from the red line indicates dynamic FRET. **(C, D)** The FRET-2CDE dynamics score is plotted against FRET efficiency. The FRET-2CDE=20 threshold is indicated as red dashed lines, and the percentage of bursts with FRET-2CDE>20 is shown in red text. **(E, F)** BVA dynamicsscores are plotted against FRET efficiency. Red diamonds show the centers of burst subgroups equally spaced along the FRET efficiency axis. The solid black lines show mean BVA-scores, and the transparent grey areas demonstrate 99.9% confidence intervals expected for static molecules, given the shot-noise present in the data. **(G, H)** The cross-correlation fFCS function is plotted against time lag. Experimental points are shown in black; the error bars were estimated by statistical bootstrapping. The fitted curve is shown in orange; the exchange times derived from the fit are highlighted with vertical red lines. The number of fluorescence bursts used for the analysis are the same as for fig. 1C: 10,167 for ZM241385, 11,961 for apo-state, 9,557 for LUF5834, 11,007 for NECA.

### FRET-2CDE and BVA confirm that agonists enhance conformational dynamics in A_2A_AR compared to apo receptor

Variations of FRET efficiency within fluorescence bursts from individual receptors suggest the presence of conformational dynamics. To analyze these variations further we used two complementary approaches: FRET-2CDE^39^ and BVA^40^. Both methods assign dynamics scores to individual molecules and are sensitive to the dynamics that are slower than the time used for FRET efficiency averaging (roughly, 100 μs for both approaches).

The FRET-2CDE score provides an unbiased way for the separation of static and dynamic subpopulations of molecules and for the comparison of their fractions in different datasets^39^. The main advantage of FRET-2CDE is that it is minimally influenced by the mean FRET efficiency in a dynamic molecule. Theoretically, static molecules should have FRET-2CDE10, while higher FRET-2CDE values should correspond to more pronounced conformational dynamics (Fig. 2C). In our data, neither the apo nor ligand-bound A_2A_AR showed a clear separation of different receptor subpopulations along the FRET-2CDE axis, but the agonists did increase the mean FRET-2CDE score of A_2A_AR compared to the apo or antagonist-bound receptors (Fig. 2D, Table S2). The fraction of receptors that exceeded FRET-2CDE>20 threshold was also higher for the agonist-bound receptors: 14±1 % for ZM241385, 15±1 % for apo, 24±2 % for LUF5834, 23±2 % for NECA. These results indicate that either the amplitude of the observed dynamics or the number of the inter-state transitions per burst increase in A_2A_AR upon agonist binding.

BVA provides a statistically robust way to test whether the observed variations of FRET efficiency exceed fluctuations expected from the shot noise and thus to prove conformational dynamics^40^. To apply BVA, we split bursts into consecutive photon-windows with n=5 photons in each (roughly 100 μs long), calculated standard deviations of the bin-wise FRET efficiencies within each burst, and plotted them against the mean FRET efficiency. The obtained BVA scores exceeded the 99.9% confidence interval expected from the shot noise under all four conditions (Fig. 2E,F), therefore, BVA confirmed that sub-millisecond conformational dynamics is already present in the apo and antagonist-bound A_2A_AR and further increased in the agonist-bound A_2A_AR.

### fFCS reveals two modes of A_2A_AR dynamics with different timescales

To estimate the timescales of A_2A_AR conformational dynamics we used the fFCS approach^41^. We used the photon arrival time and anisotropy information to split the photon stream from the double-labeled molecules *in silico* between the low-FRET (LF) and high-FRET (HF) channels (Fig. S4). Theoretically, if the LF and HF species are just two extremes of a heterogeneous ensemble of long-lived receptor states, then cross-correlation between the two channels will show only a diffusion-based sigmoidal component decreasing with the lag time (Fig. 2G). Contrarily, if the LF and HF species interconvert on the μs–ms timescales, the cross-correlation should decrease in the time lag region shorter than the state exchange time. (Fig. 2G) The cross-correlation function between the LF and HF channels (Fig. 2H, Table S3) showed two dynamics-conditioned anticorrelation terms on top of the positive diffusion-conditioned term:

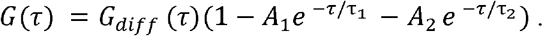

The fFCS analysis revealed fast microsecond-time (*τ_1_ = 3-20 μs*) dynamics in the aporeceptor data and in the data recorded in the presence of each of the three ligands. We believe that the fast dynamics of the dye linkers or local fluctuations of the protein can account for this term. While a single dynamic term (A_1_) described the experimental data well for the apo and the antagonist-bound A_2A_AR, the agonist-bound A_2A_AR showed an additional pronounced slower dynamics term with an exchange time of *τ_2_ = 390±80 μs*. Since the slow term appears upon agonist addition, we attribute it to the increased dynamics of the agonist-bound protein.

### PDA quantifies populations of active and inactive states in dynamic A_2A_AR

Finally, to quantify the populations of A_2A_AR in different FRET-states in the ligand-free and ligand-bound forms we used the photon distribution analysis (PDA) method^54,55^. PDA takes into account receptor dynamics and Poissonian shot noise in the measured data, and, therefore, it is preferred to a simple multi-state Gaussian fitting. For PDA, we split the fluorescence bursts into time bins of constant duration (0.5 ms, 1 ms, and 2 ms) and analyzed them globally across all apo and ligand-bound conditions. In dynamic systems, a molecule can sample several states during an individual time bin, and therefore, the FRET efficiency distribution depends on the duration of the time bin. PDA is most sensitive for picking up interconversion times on the diffusion time scale (1-10 ms); for faster or slower dynamics, PDA can be constrained *a priori* to demonstrate that the proposed model of the conformational space does not contradict the observed FRET efficiency distributions.

A model with at least three states with different FRET efficiencies was required to fit the experimental distributions. Since PDA is insensitive to fast (< 20 *μs*) dynamics observed in fFCS, the fitting models for the apo and antagonist-bound A_2A_AR did not include any interconversions between states. On the other hand, interconversion between two of the states with a fixed exchange time (*τ_2_* = 390±80 μs) as observed in fFCS was introduced in the agonist-bound PDA models. For simplicity, one state was considered long-lived and only two states were considered interconvertible. Finally, using PDA we determined the mean values and variances of inter-dye distances for each state and the populations of states under the apo and ligand-bound conditions (Fig. 3A, Fig. S5, and Table S4).

**Fig. 3.**
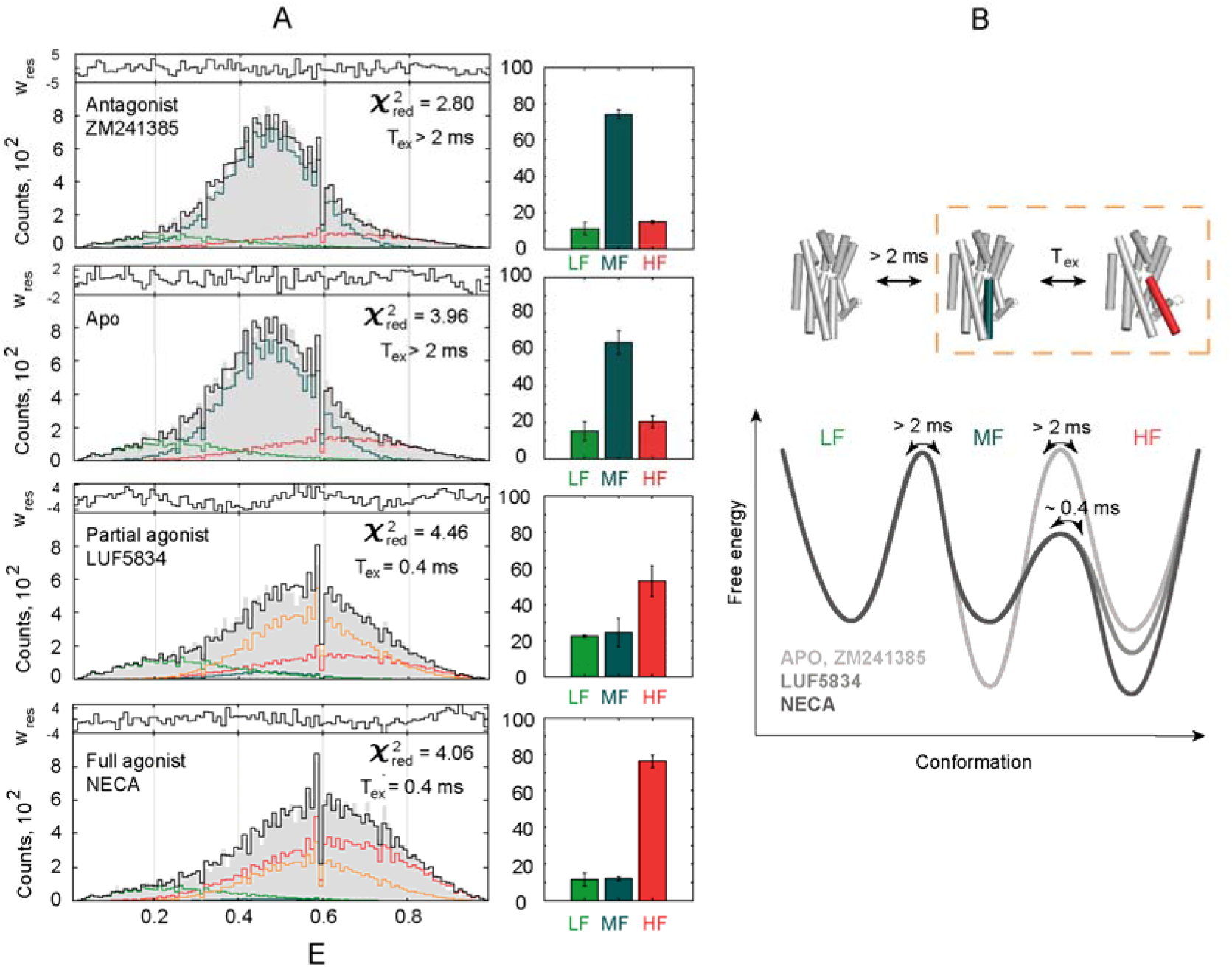
PDA quantifies parameters of the A_2A_AR three-state action model by fitting FRET efficiency distributions. **(A)** Experimental distributions of 1-ms-long time-bins derived from fluorescence bursts of double-labeled A_2A_AR (grey area) were fitted with a three state-model. The resulting fit (black line) is a sum of distributions simulated for molecules that stay in the LF (*R* = 57.9±6.0 Å, light green line), MF (*R* = 50.0±2.1 Å, dark cyan line), or HF (*R* = 45.1±4.9 Å, red line) state during the entire simulated time-bin, and the distribution for molecules that sample both MF and HF states within a time-bin (orange line). The fitting residuals are shown on the top of each panel. The bar charts on the right show relative populations of the three states, with error bars representing SD of n = 3 independent data subsets. **(B)** The three-state action model of A_2A_AR and corresponding energy landscapes for the apo and agonist-bound receptor demonstrate relative populations of the states and interstate exchange times. *τ_ex_=(k_12_ + k_21_)^-^* is the relaxation time of the exchange between the MF and HF states. TM6 is colored on the schematic (cylinder) representation of active (PDB: 5G53^59^, red) and inactive (PDB: 3EML^58^, dark cyan) structures of A_2A_AR.

The PDA results (Fig. 3A) revealed that both agonists increased the population of the highest FRET efficiency state (HF state), and decreased the population of the state with intermediate FRET efficiency (MF) state – therefore, we assume that the HF and MF states correspond to the active-like and inactive-like conformations of A_2A_AR, respectively. Approximately 10-20% of A_2A_AR molecules always stay in the low-FRET (LF) state independently of the added ligand – we speculate that these receptors are locked in a long-lived non-functional state or improperly folded. PDA converged to a model, where LF state is long-lived and MF state and HF states are interconvertible in agonist-bound A_2A_AR.

Interestingly, the active-like HF state is also observed in the apo-receptor ensemble and even in the ZM241385-bound receptors. Additionally, the sample with the full agonist NECA has a higher population of the active-like HF state compared to the partial agonist LUF5834. The small variations in state populations between the apo-receptor and the antagonist-bound receptor are below statistical significance. We discuss below the implications of these results on the basal activity, partial agonism, and inverse agonism in A_2A_AR.

## Discussion

In this study, we used smFRET to investigate the conformational dynamics of A_2A_AR. To preserve the native conformational dynamics of the receptor and minimize measurement-related artifacts we used four strategies. (1) As previous studies showed dramatic effects of commonly used detergents on GPCR conformational dynamics^56,57^ we reconstituted the receptor in nanodiscs that provide a relevant native-like lipid bilayer environment. (2) To minimize the effect of fluorophores on the receptor’s dynamics we used small organic dyes attached to strategically engineered cysteines. (3) We avoided the need to remove native cysteines and associated to that potential structural perturbations by using the previously developed in-membrane labeling procedure^49^. We showed that the mutant form of the receptor retains functional activity using the thermal shift assay and cAMP signaling assay in live HEK293T cells. (4) Finally, we studied receptors freely diffusing in solution and therefore excluded any artifacts related to their immobilization.

MFD-PIE fluorescence microscopy allowed us to filter out unlabeled and single-dye labeled receptors and to measure FRET efficiency with sub-millisecond temporal resolution in the double-labeled subpopulation of the receptors. Using various burst-wise fluorescence analysis techniques, we revealed sub-millisecond conformational dynamics in A_2A_AR. Deviation of bursts from the ‘static FRET line’ on the FRET efficiency versus donor fluorescence lifetime plot indicated nanosecond-millisecond dynamics for the apo-A_2A_AR and A_2A_AR with each of the used ligands (Fig. 2E). FRET-2CDE analysis suggested more pronounced conformational dynamics in the agonist-bound A_2A_AR than in the apo or antagonist-bound A_2A_AR (Fig. 2F). BVA confirmed that the variations of FRET efficiency among ~100 μs time-bins exceed the level expected from shot-noise (Fig. 2G). fFCS demonstrated two components in A_2A_AR dynamics: fast microsecond-time (3-20 μs) dynamics present in all samples and slower (390±80 μs) dynamics evoked with agonists (Fig. 2H). This fFCS result puts all our findings in a single self-consistent picture: both fast and slow dynamics contribute to the deviation of bursts from the ‘static FRET line’, however the fast dynamics makes almost no contributions to the FRET-2CDE scores and to the BVA distribution deviations because of their 10-fold faster timescale compared to the temporal resolution of these techniques. Meanwhile, the slower dynamics evoked with the agonists explains the increased dynamics scores in FRET-2CDE and BVA for the agonist-bound A_2A_AR. Finally, using dynamic PDA we proposed a three-state model of the A_2A_AR conformational dynamics that could fit the measured FRET efficiency histograms consistently with the fFCS findings (Fig. 3).

In our final three-state model of the A_2A_AR conformational space (Fig. 3B), the inactive-like MF and active-like HF states are not interchangeable in sub-millisecond time domain in the apo- and antagonist-bound A_2A_AR, but can interconvert on a 300-500 μs timescale in the agonist-bound A_2A_AR. The faster dynamics (3-20 μs) that contribute to conformational plasticity within these states is not sensitive to ligand binding and, therefore, likely, reflect the dynamics of the dye linkers or local fluctuations of TM6 and H8. The least populated (LF) state, presumably, corresponds to receptors locked in a long-lived non-functional state or improperly folded. Thus, our data reveal two modes of relative motions of fluorescent labels attached to residues L225C^6.27^ and Q310C^8.65^ in A_2A_AR: fast microsecond-time (3-20 μs) dynamics, present in all samples regardless of the bound ligand, and slower (300-500 μs) dynamics, evoked with agonists. In a good agreement with our findings, sub-millisecond agonist-induced conformational dynamics of the intracellular part of TM7^17^ (Y290W^7.55^) and the N-terminal end of H8^20^ (I292M^8.47^) have been shown for A_2A_AR by NMR. Although submillisecond dynamics between inactive-like conformations of TM6 (V229C^6.31^) in the apo-A_2A_AR have been reported^19^, our data suggest only limited dynamics in the ligand-free and antagonist-bound A_2A_AR.

In our experiments we could not measure slow conformational dynamics (>2 ms), because of the short residence time of individual molecules in the microscope focal spot. Our data do not indicate long-lived states in the agonist-bound A_2A_AR (besides the ligand-insensitive LF state), but the FRET-2CDE analysis shows only moderate dynamics scores, and the observed deviations from the ‘static FRET line’ can be explained by a microsecond plasticity within a long-lived conformation. Consequently, we cannot exclude that long-lived conformations can coexist with conformations that show sub-millisecond dynamics. Keeping this in mind, we nevertheless did not introduce any additional long-lived states into our final model of the A_2A_AR conformational space to avoid overfitting. Previous studies based on NMR provide complementary insights into the dynamics of long-lived (>2 ms) A_2A_AR conformations^19–21^.

The observed increase in FRET efficiency upon agonist binding was unexpected based on the available crystal structures of A_2A_AR, which predicted a decrease in the FRET efficiency, because the distance between the C’(/-atoms of the labeled residues (L225 and Q310) increases from ~40 Å in the antagonist-bound structure (PDB: 3EML^58^) to ~47 Å in the fully-active structure (PDB: 5G53^59^). Moreover, previously reported A_2A_AR-based FRET-sensors with fluorescent proteins at ICL3 and C-terminus showed a decrease in the FRET efficiency upon receptor activation^42,43^. To explain this discrepancy, we proposed that the dyes are not randomly distributed (Fig. 1B), but rather occupy preferred locations within the volume accessible with their ~15 Å linkers. To test this hypothesis we performed 1-μs long molecular dynamics (MD) simulations, which revealed that the dye attached to TM6 might preferentially locate between the intracellular tips of TM3 and TM5 in the inactive conformation and enter the G-protein-binding cavity of the receptor in the active conformation (Fig S8). These preferred conformations of the dye would result in a significant decrease of the mean inter-dye distance upon receptor activation (from 5 nm to 3 nm), which would in turn lead to an increase in the mean FRET efficiency. Thus, our MD simulations provide a plausible explanation for the observed increase in the FRET efficiency upon A_2A_AR activation.

The PDA analysis of our data suggests that the partial agonist LUF5834 and the full agonist NECA stabilize 53±8 % and 76±3 % of A_2A_AR molecules, respectively, in the same active-like HF conformation. A similar mechanism for partial agonism in A_2A_AR has recently been demonstrated via NMR with isotope-labeled methionine residues located in different structural domains (I106M^3.54^, M140^4.61^, M211^5.72^, and I292M^8.47^) of the receptor^20^. On the other hand, two other NMR-based studies have suggested that LUF5834 either stabilizes a distinct, not a fully active conformation^19^, or has no effect on the A_2A_AR conformation^21^. Our data do not support the existence of a separate partially active conformation of A_2A_AR stabilized with LUF5834 that would be distinct from the fully active conformation stabilized with NECA. On the other hand, we cannot exclude that such partially active state could not be resolved in our data because small differences in FRET efficiency, the high photon shotnoise in single-molecule experiments, or the broadening of the FRET-distribution due to variations of photophysical parameters of the dyes.

Additionally, we observed that 20±3 % of apo A_2A_ARs exhibit an active-like conformation, which could translate into a moderate basal activity of the receptor. Two previous NMR-based studies have addressed the molecular mechanisms of A_2A_AR basal activity. One study reported a 70% population of pre-active and fully active states in the apo-ensemble^19^. Another study has reported negligible basal activity and showed that in the agonist-bound A_2A_AR unique previously unpopulated conformations emerge^21^. A recent review suggests that discrepancies between these two works could arise from differences in used constructs, ^19^F reporters, their attachment sites, or in selected membrane mimicking systems (MNG/CHS versus DDM/CHS micelles)^60^. The contradictory estimations of the basal activity of A_2A_AR should be put in the context of a similar heterogeneity of results provided by cell-based signaling assays. In different experiments, the basal activity of A_2A_AR was reported to reach from 0-20%^58,61–63^, to 20-40%^43,64,65^ or even 40-70%^66–69^ Cell assays are affected by different A_2A_AR expression levels and cell lines used^62,68^. It has been shown that the C-terminal truncation of A_2A_AR impairs its basal activity - this can play an important role for our study as well as previous NMR-based works^64^.

Finally, our measurements show that ZM241385 does not change the distribution of FRET efficiency compared to apo conditions and therefore we do not observe inverse agonism of ZM241385. Because many studies reported negligible basal activity of A_2A_AR, ZM241385 is widely referred to as A_2A_AR antagonist^11,58,70^. The recent ^19^F NMR study, where no basal activity was detected for A_2A_AR, correspondingly did not register any conformational changes induced with ZM241385^21^. On the other hand, those works that identified significant basal activity of A_2A_AR frequently reported inverse agonism of ZM241385^43,62,63,65,68,71^. In line with these findings, the ^19^F-NMR study that has reported 70% basal activity also showed inverse agonism of ZM241385^19^. Notably, it was previously shown that ZM241385 can lose inverse agonist activity if tested not in cells, but in isolated membranes^64^. This latter result suggests that intracellular interaction partners can play an important role in both basal activity and inverse agonism, explaining both heterogeneity in published functional data and our results.

The multi-state conformational behavior of GPCRs delineates their complex pharmacology and, therefore, challenges modern drug design. We believe that new methods showing how GPCR activity is modulated on a molecular level will facilitate the design and discovery of drugs with novel beneficial properties. Here we demonstrated a novel strategy to observe conformational dynamics of a GPCR in solution, yet in a close-to-physiological environment of lipid nanodiscs using intramolecular smFRET measured via the MFD-PIE approach. Our measurements combined fluorescence intensity, lifetime, and anisotropy information to characterize the sub-millisecond conformational dynamics of TM6 and H8 in A_2A_AR and shed light on molecular mechanisms of basal activity and partial agonism in the receptor. The general strategy developed in our work can be extended to study the effects of various modulators (ligands, ions, lipids, etc.), membrane-mimicking systems (micelles, lipid nanodiscs, liposomes, etc.) and genetic modifications on the activity of A_2A_AR and, in perspective, other GPCRs.

## Methods

### Protein expression, purification and labelling

The gene encoding the human A_2A_ adenosine receptor (1-316 aa) (UniProt C9JQD8) was synthesized *de novo* (Eurofins). The nucleotide sequence was optimized for *Leishmania tarentolae* expression with the GeneOptimizer software (ThermoFisher Scientific). KpnI restriction site was introduced at the C-terminus and used for polyhistidine tag (H9) fusion. The final construct was cloned into the integrative inducible expression vector pLEXSY_I-blecherry3 (Jena Bioscience, Germany) via the BglII and NotI restriction sites. L225C^6.27^ and Q310C^8.65^ mutations were introduced by PCR.

*Leishmania tarentolae* cells of the strain LEXSY host T7-TR (Jena Bioscience) were transformed with the A_2A_AR expression plasmids linearized by the SmiI restriction enzyme. After the clonal selection, the transformed cells were grown at 26 °C in the dark in shaking baffled flasks in the Brain-Heart-Infusion Broth (Carl Roth, Germany) supplemented with 5 μg/mL Hemin (AppliChem), 50 U/mL penicillin and 50 μg/mL streptomycin (both antibiotics from AppliChem). When OD_600_=1 was reached, 10 μg/mL tetracycline was added, and incubation continued for additional 24h.

The harvested cells were disrupted in an M-110P Lab Homogenizer (Microfluidics) at 10,000 psi in a buffer containing 50 mM NaH_2_PO_4_/Na_2_HPO_4_, pH 7.6, 0.2 M NaCl, 20 mM KCl, 10 mM MgCl_2_, 10% glycerol (w/v), 1 mM EDTA, 2 mM 6-aminohexanoic acid (AppliChem), 50 mg/L DNase I (Sigma-Aldrich) and cOmplete protease inhibitor cocktail (Roche). The membrane fraction of the cell lysate was isolated by ultracentrifugation at 120,000 g for 1 h at 4 °C. The pellet was resuspended in the same buffer but without DNase I and stirred for 1h at 4 °C. The ultracentrifugation step was repeated again.

Finally, the membranes were resuspended in the labelling buffer containing 50 mM HEPES, pH 7.0 10 mM MgCl_2_, 20 mM KCl, 2 mM 6-aminohexanoic acid, and cOmplete and mixed with Atto643 maleimide (ATTO-TEC) and Alexa488 maleimide (Invitrogen), dissolved in dimethyl sulfoxide (0.5 mg of each fluorescent label per 10 g of cells). Labeling reactions were carried out overnight in the dark at 4 °C on a roller mixer.

The next day, membrane fractions were pelleted by ultracentrifugation at 120,000 g for 1 h at 4 °C and washed twice with the labelling buffer for removal of unbound fluorescent labels. For solubilization, membranes were resuspended in a buffer containing 20 mM HEPES, pH 8.0, 800 mM NaCl, 5 mM MgCl_2_, 10 mM KCl, 2 mM 6-aminohexanoic acid, cOmplete with 4 mM theophylline (Sigma-Aldrich) and 1% n-Dodecyl β-maltoside (DDM) (Glycon Biochemicals) / 0.2% cholesteryl hemisuccinate (CHS) (Merck) (w/v) and left on the stirrer for 2 h at 4 °C in the dark. The insoluble fractions were removed by ultracentrifugation at 120,000 g for 1 h at 4 °C. The supernatants were loaded on an Ni-NTA resin (Cube Biotech) and incubated in the batch-mode overnight in the dark at 4 °C.

The next morning, proteins bound to Ni-NTA resin were washed with 10 column volumes of the first washing buffer: 50 mM HEPES, pH7.5, 800 mM NaCl, 25 mM imidazole, 10 mM MgCl_2_, 8 mM ATP (Sigma-Aldrich), 2 mM 6-aminohexanoic acid, 0.1 mM phenylmethylsulfonyl fluoride, 4 mM theophylline, cOmplete, 0.1% DDM / 0.02% CHS. Then, columns were washed with 10 column volumes of the second washing buffer: 50 mM HEPES, pH 7.5, 800 mM NaCl, 50 mM imidazole, 2 mM 6-aminohexanoic acid, 0.1 mM phenylmethylsulfonyl fluoride, 4 mM theophylline, cOmplete, 0.1% DDM / 0.02% CHS (w/v). Finally, proteins were eluted with 5 column volumes of the elution buffer: 25 mM HEPES, pH 7.5, 800 mM NaCl, 220 mM imidazole, 2 mM 6-aminohexanoic acid, 0.1 mM phenylmethylsulfonyl fluoride, cOmplete, 0.1% DDM / 0.02% CHS (w/v). The eluates were subjected to size-exclusion chromatography on a Superdex 200 Increase 10/300 GL column (GE Healthcare Life Sciences) in a buffer containing 20 mM HEPES, pH 7.5, 800 mM NaCl, 1 mM EDTA, 2 mM 6-aminohexanoic acid, cOmplete, 0.05% DDM / 0.01% CHS (w/v). Fractions, corresponding to A_2A_AR monomers, were pulled and subjected to nanodisc reconstitution.

### Nanodisc reconstitution

Membrane Scaffold Protein 1D1 (MSP1D1) was expressed in *E.coli* using gene with an N-terminal 6XHis-tag and up stream TEV-protease site cloned into pET28a(+) (Addgene plasmid #20061^50^). MSP1D1 was purified using IMAC^72^ with further cleavage of 6xHis-tag by TEV protease (Sigma-Aldrich). The lipid mixture of 1-palmitoyl-2-oleoyl-sn-glycero-3-phosphocholine (POPC): 1-palmitoyl-2-oleoyl-sn-glycero-3-phospho-(1’-rac-glycerol) (POPG) (Avanti Polar Lipids) in chloroform was prepared at a molar ratio 7:3. The lipid film was dried under a gentle nitrogen stream, followed by removal of the solvent traces under vacuum, and then solubilized in 200 mM sodium cholate. The purified A_2A_AR in DDM/CHS micelles was mixed with MSP1D1 and the POPC:POPG lipids at a molar ratio A_2A_AR:MSP1D1:lipids=0.2:1:60. The final sodium cholate concentration was adjusted to 20 mM, the typical final receptor concentration was 0.1 mg/mL. After 1 h incubation at 4 °C, the mixture was incubated with wet Bio-Beads SM-2 (Bio-Rad, 0.4 g of beads for 1 mL reaction, beads were washed in methanol and equilibrated with 20 mM HEPES, pH 7.5, 800 mM NaCl, 1 mM EDTA) overnight at 4 °C in the dark. The next morning, the beads were discarded and the supernatant was supplemented with a fresh portion of Bio-Beads for an additional 4 h incubation. Finally, A_2A_AR reconstituted into nanodiscs was subjected to size-exclusion chromatography on a Superdex 200 Increase 10/300 GL column (GE Healthcare) in a buffer containing 20 mM HEPES, pH 7.5, 150 mM NaCl, 1 mM EDTA, 2 mM 6-aminohexanoic acid, cOmplete. Fractions containing labeled receptors were combined together and used for further experiments.

### Thermal shift assay

To show that the A_2A_AR mutant (L225C^6.27^/Q310C^8.65^) retains ligand-binding activity in lipid nanodiscs, we used the fluorescent thermal stability assay^51^. The studies were carried out on a Rotor-Gene Q 6 plex (QIAGEN) instrument at a heating rate of 2 °C/min and a temperature range of 25-90 °C. The excitation wavelength was set at 387 nm and the emission wavelength was 463 nm. The A_2A_AR concentration was about 2 μM. Buffer conditions: 20 mM HEPES, 150 mM NaCl, 1 mM EDTA, 2 mM 6-aminohexanoic acid, pH 7.5. To obtain a good fluorescent intensity we used a 2.5-fold molar excess of CPM dye (7-Diethylamino-3-(4’-Maleimidylphenyl)-4-Methylcoumarin, Invitrogen) to protein. To prepare protein for the ligand-binding measurements we added 200 μM of ZM241385 or NECA and incubated for 1h in the dark at +4 °C. The thermal denaturation assay was performed in a total volume of 50 μL (Fig. S1E).

### Measurement of A_2A_AR surface expression and Gs-signaling

For A_2A_AR functional assays, the A_2A_AR (WT or Q310C^8.65^/L225C^6.27^ mutant) gene (GenScript) was optimized for eukaryotic expression with an N-terminal hemagglutinin signal sequence (MKTIIALSYIFCLVFA) followed by the FLAG tag epitope (DYKDDDDK) and C-terminal 10×His tag were cloned into pcDNA3.1(-) at BamHI(5’) and HindIII(3’). The surface expression of A_2A_AR was determined by the whole-cell ELISA assay^73^. Briefly, HEK293FT cells were seeded in a 100 mm cell culture plate and transfected separately with 10 μg of each expression plasmid DNA (pcDNA3.1(-)_A_2A_AR(WT), pcDNA3.1(-)_A_2A_AR(Q310C^8.65^/L225C^6.27^) or pCDNA3.1(-) as a negative control) using a common Lipofectamine 3000 protocol. The plates were incubated for additional 12–18 h at 37 °C, 5% CO_2_. The HRP-conjugated anti-FLAG M2 antibody (A8592, Sigma) at a dilution of 1:2000 in TBS with 1% protease-free BSA (A3059, Sigma) and TMB ready-to-use substrate (T0565, Sigma) were used for the ELISA procedure. For normalization on cells quantity Janus Green B (Sigma) staining was used, and the absorbance ratio A_450_/A_595_ was calculated. Measurements were performed in triplicate for WT and mutant A_2A_AR as well as for empty-vector transfected cells. Measured values of A_450_/A_595_ were normalized so that the mean expression level of WT A_2A_AR was 100% (F_WT_ = 100±6 %, SDs for n=3 measurements are given). The double mutant form of the receptor showed only slightly lower expression level than WT: F_L225C/Q310C_ = 73±7 %. Empty-vector transfected cells showed only marginal anti-FLAG antibody binding: F_EV_ = 1±1 %.

For evaluation of the A_2A_AR signaling activity, we checked the effect of the agonist NECA on cAMP responses in transfected cells. For cAMP determination, we used the Bioluminescence Resonance Energy Transfer (BRET) approach with the EPAC biosensor^74^. The cAMP BRET biosensor was kindly provided by professor Raul Gainetdinov^75^. Transfections were carried out with Lipofectamine 3000 (Thermo) using HEK293T cells seeded in a 100 mM cell culture plate, receptor cDNA vectors pcDNA3.1(-)_A_2A_AR(WT), pcDNA3.1(-)_A_2A_AR(Q310C^8.65^/L225C^6.27^) (10 μg each) and the EPAC biosensor cDNA vector (1 μg) needed for evaluation of the cAMP production. Transfected cells were split into 96-well plates at 10^5^ cells per well. On the following day, 70 μL of PBS were added to each well followed by addition of 10 μL of a 50 μM coelenterazine-h solution (Promega). After 10-min incubation, either 10 μL of buffer or 10 μL of NEC A at different concentrations in PBS were added, and the plate was then placed into a CLARIOstar reader (BMG LABTECH, Germany) with a special BRET filter pair (475±30 nm _ coelenterazine-h and 530±30 nm _ YFP). The BRET signal was calculated as the ratio of the light emitted at 530 nm to the light emitted at 480 nm. Three independent experiments with three technical replicas in each were conducted. For pEC_50_ evaluation, dose-response curves from three technical replicas were averaged and analyzed. Mean and S.D. of pEC_50_ among three biological samples were calculated (Fig. S1F).

### Confocal MFD-PIE setup

For single-molecule experiments, a home-built multi-parameter fluorescence detection microscope with pulsed interleaved excitation (MFD-PIE)^36^ was used (see scheme of the setup in Fig. S2). Two lasers were used: a pulsed 483-nm laser diode (LDH-P-C-470, Picoquant) and a pulsed 635-nm laser diode (LDH-P-C-635B, Picoquant), with alternating at 26.67 MHz pulses, delayed by 18 ns with respect to each other. Sample emission was transmitted through a pinhole and spectrally split. Both, the blue range and red range were split by polarization into two detection channels. Photons were detected by four avalanche photodiodes (PerkinElmer or EG&G SPCM-AQR12/14, or Laser Components COUNT BLUE): B_║_ (blue-parallel), B_⊥_ (blue-perpendicular), R_║_ (red-parallel) and R_⊥_ (red-perpendicular) (Fig. S2), which were connected to a TCSPC device (SPC-630, Becker & Hickl GmbH). Microscope alignment (excitation light guiding, objective lens correction collar, pinhole, detectors) was done using real-time fluorescence correlation spectroscopy (FCS) on freely diffusing Atto488-COOH and Atto655-COOH in water. For more details about the used equipment the reader is referred to ref.^76^

### smFRET data recording

Samples of double-labelled A_2A_AR in nanodiscs were diluted in a buffer, containing 20 mM HEPES, pH 7.5, 150 mM NaCl, 1 mM EDTA, 2 mM 6-aminohexanoic acid to a protein concentration of 0.5-2 nM. To measure the effects of ligand binding, samples were supplemented with either 10 μM ZM241385, 10 μM LUF5834 or 10 μM NECA and incubated for 30 min at +4 °C. After the incubation, the samples were transferred to a Nunc Lab-Tek Chambered coverglass (Thermo). smFRET experiments were performed at 100 μW of 483 nm and 50 μW of 635 nm excitation. Measurements were recorded at room temperature (22 °C), samples were replenished every 30 min. With all filters applied (see *Selection of double-labeled, donor-only and acceptor-only subpopulations*), 9,000 – 12,000 bursts corresponding to double-labeled molecules were collected for each sample: 11,961 for apo, 10,167 burst for ZM241385, 9,557 for LUF5834, and 11,007 for NECA. Background scattering information was obtained via a buffer measurement under identical condition.

### Software

All simulations and analyses of experimental data were performed in the software package PAM (PIE Analysis with MATLAB)^77^. The software is available as a source code, requiring MATLAB to run, or as pre-compiled standalone distributions for Windows or MacOS at http://www.cup.uni-muenchen.de/pc/lamb/software/pam.html and hosted in Git repositories under http://www.gitlab.com/PAM-PIE/PAM and http://www.gitlab.com/PAM-PIE/PAMcompiled. A detailed manual is located under http://pam.readthedocs.io. Details of smFRET data treatment are given in Supplementary methods.

## Supporting information

Supplementary text 1. Topology for Atto647N-maleimide.

Supplementary text 2. Topology for Alexa488-C5-maleimide.

## Acknowledgements

This work is supported by RFBR research grant 20-34-70034. VC acknowledges that the University of Southern California is his primary affiliation. A.L., V.G., A.M. and V.B. are thankful for the Ministry of Science and Higher Education of the Russian Federation (agreement #075-00337-20-03, project FSMG-2020-0003).

## Author contributions

OV performed receptor expression. IM and OV performed receptor labeling, purification, and nanodisc reconstitution. TG and IM performed ensemble-TCSPC measurements. IM and QC measured fluorescence spectra. QC and JHe prepared the instrumentation for smFRET experiments. IM performed smFRET experiments, analyzed smFRET data and drafted the manuscript. JHe, ABa, SW, ABo and VB contributed to the analysis of smFRET data. AGe performed cell functional assay. PKh performed TSA functional assay with the contribution of AM, AL, AGu and PKu. PO performed molecular dynamics simulations. IM, VB, JHe, VC, TG, ABo, AGu, AM, PKh, VG discussed the data, analysis, and contributed to writing the manuscript. IM, VB, JHe, VC, TG, JHo conceived the study. All authors commented on and edited the manuscript. VB, and JHe supervised the project.

## Competing interests

Authors declare no competing interests.

## Notes

### Competing Interest Statement

The authors have declared no competing interest.

