## Supplementary Information for "Sub-millisecond conformational dynamics of the A_2A_ adenosine receptor revealed by single-molecule FRET"

Maslov et al.

### Supplementary methods

#### Fluorescence spectra characterization

For fluorescence spectra characterization, diluted (<5 µM) apo-A_2A_AR samples were placed in a quartz cuvette (10 mm path length). Excitation and emission spectra were recorded using an Edinburgh Instruments FLS980 spectrometer corrected for the wavelength-dependent throughput and sensitivity of the detector. Fluorescence in the acceptor’s emission spectral range after irradiation in the donor’s excitation spectral range indicated FRET in the double-labeled receptor samples (Fig. S1C).

#### Ensemble-based fluorescence lifetime measurements

The time-resolved detection of the fluorescence decay of Apo-A_2A_AR labeled with Alexa488 and Atto643 was performed with a Fluotime100 fluorescence spectrophotometer (Picoquant, Berlin, Germany) based on a picoHarp300 unit and using a pulsed diode laser (LDH-440; center wavelength 440 nm; pulse width: 54 ps; repetition frequency: 10 MHz) as an excitation source. Fluorescence decay curves were measured at 665 nm under magic angle conditions by time-correlated single-photon counting (TCSPC) allowing to determine fluorescence lifetimes down to 100 ps^1^. Decay curves were analyzed by iterative reconvolution of the instrument response function, IRF(t), with an exponential model function, M(t), using the FluoFit software (version 4.4; Picoquant).

Fitting the measured TCSPC-delay signal with a monoexponential decay (Fig. S1D) did not allow a satisfactory description of the acceptor fluorescence intensity time trace, while a biexponential fit was sufficient and yielded two components: one with a positive amplitude (normal fluorescence decay) and one with a negative amplitude (rise term). The rising term is expected for FRET and cannot originate exclusively from the direct excitation of Atto 643 with a 440-nm laser. Therefore, fluorescence lifetime measurements of labeled mutant protein in bulk solution also confirmed that there is a fraction of double-labeled receptors that exhibit FRET in the sample.

#### Burst Identification

For single-molecule data, we employed a two-color MFD all-photon burst search algorithm^2^ using a 500 µs sliding time window (min. 50 photons per burst, min. 5 photons per time window). A 0-20-ms burst duration cutoff was applied to remove sparse (< 1%) slow moving aggregates.

#### FRET efficiency and Stoichiometry

The absolute burst-averaged FRET efficiency E was calculated with:

$E =\frac{F_{BR}- ct* F_{BB}-de* F_{RR}}{\gamma F_{BB}+F_{BR}- ct* F_{BB}-de* F_{RR}}$

where *F_BR_* = *S_BR_* – *B_BR_* is the background-corrected number of photons in the red detection channels independently of the polarization after blue excitation (with *S_BR_* and *B_BR_* being the summed intensity and background, respectively, in the time gates *BR_∥_* and *BR_⊥_*); *F_BB_* = *S_BB_* – *B_BB_* is the background-corrected number of photons in the blue detection channels after blue excitation (with *S_BB_* and *B_BB_* being the summed intensity and background, respectively, in the time gates *BB_∥_* and *BB_⊥_*), *F_RR_* = *S_RR_* – *B_RR_* is the background-corrected number of photons in the red detection channels after red excitation (with *S_RR_* and *B_RR_* being the summed intensity and background, respectively, in the time gates *RR_∥_* and *RR_⊥_*), *de* – the correction factor for direct excitation of the acceptor with the 483 nm laser, *ct* – the correction factor for the emission crosstalk of the donor in the acceptor channel, and *γ* – the relative detection efficiency of the donor and acceptor^3^.

The corrected stoichiometry ratio *S* was calculated with:

$S = \frac{\gamma F_{BB}+F_{BR}- ct* F_{BB}-de* F_{RR}}{\gamma F_{BB}+F_{BR}- ct* F_{BB}-de* F_{RR}+\beta F_{RR}}$,

where *ꞵ*-factor accounts for different detection efficiencies of the donor and the acceptor.

#### Correction factors

For correction, first, the background was subtracted from the experimental signals. Then, the donor emission crosstalk (*ct* =  0.0059) and acceptor direct excitation (*de*  =  0.024) factors were determined directly from the measurements and applied to correct the data ^3^. For correction purposes, we preliminarily (see the final selection criteria for other analyses in the section “Selection of double-labeled, donor-only and acceptor-only subpopulations”) selected double-labeled molecules using the kernel-density estimator (ALEX-2CDE<15)^4^, FRET efficiency (0.1<*E*<1), and stoichiometry (0.2<*S*<0.6), corrected for channel crosstalk (*ct*) and direct excitation (*de*). For these selected molecules, *E* was plotted vs 1/*S*, and a straight line was fitted to obtain the correction factors:

$\gamma= \frac{\Omega-1}{\Omega+\Sigma-1}$

$\beta=\Omega+\Sigma-1$,

where $\Omega$ is the intercept and $\Sigma$ is the slope of the fit. Finally, *γ* = 0.69 and *ꞵ*  =  1.9 were obtained.

#### Burst-wise fluorescence lifetime

To estimate the single-molecule burst-averaged fluorescence lifetimes of the donor (τ_D_) and acceptor (τ_A_), the maximum likelihood estimator approach was used^5^.

#### Burst-wise steady-state fluorescence anisotropies

Burst-wise steady-state fluorescence anisotropies of the donor (r_D_) and the acceptor (r_A_) were calculated from the respective fluorescence intensities:

$r= \frac{G F_{||}-F_{\perp}}{G F_{||}+2F_{\perp}}$

where *G* is the correction factor for different detection efficiencies in the two polarization channels (*G_B_*  =  0.99, *G_R_*  =  1.13), *F_∥_* is the intensity in the time gate BB_∥_ (donor) or RR_∥_(acceptor), and F_⊥_ is the intensity in the time gate BB_⊥_ (donor) or RR_⊥_ (acceptor).

#### Selection of double-labeled, donor-only and acceptor-only subpopulations

To select single-labeled or double-labeled subpopulations of molecules, we used specific restrictions for the stoichiometry *S*, FRET efficiency *E*, and kernel-density estimator ALEX-2CDE, as shown below. The data were further cleaned using the following fluorescence lifetime and anisotropy filters.

Donor-only molecules:$ALEX-2CDE>20$,$-0.1<E<0.1$, $0.9<S<1.1$,$0.1 ns<\tau_{D}<6 ns$, $-0.2<r_{D}<0.6$.

Acceptor-only molecules: $ALEX-2CDE>20$,$0.6<E<1.1$, $-0.1<S<0.2$, $0.1 ns < \tau_{A}< 8 ns$, $-0.2<r_{A}<0.6$.

Double-labeled molecules: $ALEX-2CDE<15$,$0.1<E<1.0$,$0.2<S<0.8$, $0.1 ns<\tau_{D}<4.5 ns$, $0.1 ns<\tau_{A}<8 ns$, $-0.2<r_{D}<0.6$, $-0.2<r_{A}<0.6$.

#### Photon distribution analysis (PDA)

Static PDA was carried out to obtain the absolute inter-dye distance distribution as previously described^6^, assuming three Gaussian-distributed states. Dynamic PDA with one static and two interconverting dynamic states was carried out to account for the conformational dynamics revealed by other analysis approaches^7^. Practically, for each smFRET dataset, raw bursts were re-binned in different time bins (0.5, 1, and 2 ms), and three histograms were constructed and analyzed simultaneously. Only bins with at least 20 and maximally 300 photons (to reduce calculation time) were further analyzed using PDA. Bins with uncorrected stoichiometry S_PR_ below 0.2 or above 0.6 were removed from the analysis, because of suspected complex acceptor photophysics or photobleaching. A three-state model with Gaussian distance distributions was used to generate a library of simulated FRET efficiency values, which was subsequently fitted to the experimental corrected FRET efficiency histogram using a reduced χ^2^ –guided simplex search algorithm. Correction parameters *γ* = 0.69, *ct*  =  0.0059 and *de*  =  0.024, as well as the average background count rates in the donor and the acceptor channels after donor excitation were used to calculate the corrected FRET efficiency for PDA. The mean and width of all Gaussian distributed sub-states were globally optimized over all (three ligand and apo) conditions. State areas A_i_ and interconversion rates constants k_12_ and k_21_ were globally optimized for each sample. An fFCS-guided PDA fit was performed with a fixed exchange time *T_ex_  =  (k_12_+k_21_)^-1^* and *k_12_/k_21_* ratio optimized globally over a single sample. The exchange time was fixed to values determined from fFCS (τ_2_) for agonists, and to virtual infinity (>1,000 ms) for the antagonist-bound or ligand-free receptors (to account for reduced amplitude of dynamic term). Criteria for a good fit were a low (< 4) global reduced- χ^2^ value. The resulting parameters are presented as the means ± SD of three independent experiments in Fig. S5, Table S4.

Besides the fFCS-constrained dynamic PDA, we also tested a three-state static PDA (Fig. S6, Table S5) as well as an unconstrained dynamic PDA (Fig. S6, Table S6). The comparison between the static and dynamic analyses showed that PDA in our case is not robust enough to distinguish between the dynamic and static models. We considered the fFCS-constrained analysis as the best option, because it takes into consideration the insights into A_2A_AR dynamics obtained from other data analysis modes and thus provides a basis for a self-consistent action model of A_2A_AR (Fig. 3B). Notably, the fFCS-constrained PDA has the same number of fitting parameters as the simplest, static PDA, because the only additional parameter (exchange time) is determined in fFCS and fixed for PDA.

#### Burst Variance Analysis (BVA)

BVA was performed as described^8^. Each fluorescence burst was segmented into M_i_ bins of n = 5 consecutive photons; the proximity ratio ε_ij_ was calculated for each bin by the ratio N_a_/n, where N_a_ is a number of acceptor photons within the bin. From the resulting set {ε_ij_} and the burst-wise proximity ratio PR_i_, the standard deviation is estimated as:

$s_{i} = \sqrt{\frac{1}{M_{i}}\sum_{j=1}^{j=M_{i}} \left( \epsilon_{ij}-PR_{i} \right)^{2}}$.

The burst-wise *s_i_* values were plotted against burst-wise FRET efficiency.

Bursts were grouped in N = 20 equally-spaced intervals by the burst-wise proximity ratio *PR_i_*; only groups with >100 bursts were analyzed. Within each group, the mean value of {ε_ij_} was determined, and the corresponding FRET efficiency value was calculated using correction factors *ct*, *de*, *γ*, and *ꞵ* (eq. 29 from ^9^):

$$E = \frac{1-(1+ct+\gamma\beta\cdot de)(1- PR)}{1 - (1+ct-\gamma)(1 - PR)}$$

The standard deviation of {ε_ij_} within each group was plotted against FRET efficiency.

For comparison, the theoretical ‘static’ standard deviation *s* was determined:

$s = \sqrt{\frac{PR (1-PR)}{n}}$.

The 99.9% confidence interval for *s* was determined from simulated ‘static’ bursts, given the same number of bursts in each group. The theoretical ‘static’ standard deviation and confidence intervals were plotted against corrected FRET efficiency (Fig. 2D).

#### Filtered Fluorescence Correlation Spectroscopy (fFCS)

The mathematical background of fFCS was described in detail^10^. We built two reference TCSPC patterns corresponding to the ‘low-FRET’ pseudo-species (*p_j_^LF^*) and ‘high-FRET’ pseudo-species (*p_j_^HF^)* (Fig. S4)*.* For this, we merged all four smFRET datasets with different ligand conditions; bursts corresponding to double-labeled receptors with *E<0.3* were used to build *p_j_^LF^*, bursts corresponding to double-labeled receptors with *E>0.7* were used to build *p_j_^HF^.* TCSPC channels for BB_∥_ ,BB*_⊥_,* BR_∥_, and BR*_⊥_* excitation and emission channels were stacked into a single array and indexed with *j* for global analysis. Using the reference TCSPC patterns, *p_j_^LF^* and *p_j_^HF^* filters *f_j_^LF^* and *f_j_^HF^* were calculated as described^10^. To reduce noise in fFCS filters, at this step TCSPC bin was increased to 100 μs.

Using the reference filters *f_j_^LF^* and *f_j_^HF^* and the fluorescence signal *S_j_*, the correlation function *G(τ)* was calculated for each dataset:

$G(\tau)^{(i,m)} = \frac{<(\sum_{j=1}^{C} {f_{j}}^{(i)}S_{j}(t))\cdot(\sum_{j=1}^{C} {f_{j}}^{(m)}S_{j}(t))>}{<(\sum_{j=1}^{C} {f_{j}}^{(i)}S_{j}(t))>\cdot<(\sum_{j=1}^{C} {f_{j}}^{(m)}S_{j}(t))>}-1$.

Only bursts from double-labeled molecules were taken into account; a 10 ms time-window was introduced to reduce artifacts related to the burst-wise FCS analysis.

The cross-correlation functions (G^LF,HF^ or G^HF,LF^) were fit using equation:

$G^{\left( i,m \right)}\left( \tau\right)= {G^{\left( i,m \right)}}_{diff}\left( \tau\right)\left( 1 -{A_{1}}^{\left( i,m \right)}e^{-\frac{\tau}{\tau_{1}}}-{{A_{2}}^{\left( i,m \right)}}e^{-\frac{\tau}{\tau_{2}}} \right)$,

where the diffusion-limited term is:

${G_{diff}}^{(i,m)}(\tau) =\frac{1}{\sqrt{8}N^{(i,m)}} \frac{1}{(1+\tau/T_{D})(1+\tau/{p^{2}T}_{D})^{1/2}}$.

First, we fitted cross-correlation curves for the apo and ligand-bound conditions without any additional constrains (Table S3A). This preliminary fitting showed that the amplitude A_2_ was higher for the agonist-bound receptors than for the apo or antagonist-bound receptors. Finally, we found that a simpler fitting model with A_2_  =  0 for the apo or antagonist-bound receptors, and T_D_ and T_2_ set global across all four experimental conditions, allowed a satisfactory fitting of the experimental curves (Table S3B). The resulting cross-correlation curves were normalized using N^(i,m)^ and offset and plotted in Fig. 2H.

#### Molecular dynamics simulations

The initial model of the A_2A_AR in the inactive state (amino acids 3-316) embedded in membrane was prepared using the CHARMM-GUI web-service^11^ based on the structure of a thermostabilized A_2A_AR in complex with ZM241385 (PDB ID: 3PWH)^12^. The thermostabilized mutations were mutated back to native amino acids and the missing regions were added using MODELLER^13^ with an exception of the loop 212-223, which was omitted to prevent possible interference with the fluorescent label at the position 225 and thus improve its sampling. The structure of A_2A_AR in complex with mini-Gs (PDB ID 5G53)^14^ was used as a template to model the C-terminus of H8 missing in 3PWH. The Atto647N-maleimide and Alexa488-C5-maleimide fluorescent labels were attached at the positions 225 and 310 by aligning the backbone atoms of the modified cysteine residues with the bound fluorescent labels to the backbone atoms of corresponding residues of the protein. Two simulations were performed: one for each double-labeled variant of A_2A_AR. In these simulations, we used the Atto647N-maleimide dye instead of its derivative Atto643-maleimide used in the experiment, because the structure of the latter was not published. The resulting solvated system contained 83,191 atoms including 59 POPG lipids, 177 POPE lipids, 123 sodium, and 72 chloride ions. The simulation box had total dimensions of 9.10 × 9.10 × 9.62 nm^3^. All ionizable amino acids were modeled in their standard ionization state at pH 7. The CHARMM-GUI recommended protocol was applied for the initial energy minimization and equilibration of the system. During all of the equilibration steps, the force constants of the harmonic positional restraints on lipids were gradually reduced to zero while those on the protein Cα-atoms were left intact.

The equilibration simulations were followed by targeted MD simulation^15^ in order to steer the system to the fully active state while inducing minimal effects on the system. The 5G53 structure was used as a target for targeted simulations to the fully active state. For the targeted MD simulations, the Nose–Hoover thermostat and the Parrinello–Rahman barostat were used. The temperature and pressure were set to 313.3 K and 1 bar with temperature and pressure coupling time constants of 1.0 ps^−1^ and 0.5 ps^−1^, respectively. Each targeted simulation was run for 100 ns with the force constant of 50,000 kJ/mol applied to the protein Cα-atoms only.

The production simulations for the inactive and active states were run for 1,000 ns in NVT ensemble (maintained by the Nose–Hoover thermostat with T_ref_ = 313.3 K, temperature coupling time = 1.0 ps^−1^) with the protein Cα-atoms and all heavy atoms of lipids constrained by harmonic potentials (1,000 kJ/mol/nm^2^). The fluorescent labels were coupled separately to the heat bath (T_ref_ = 450 K), what was shown previously to enhance conformational sampling^16^.

All MD simulations were performed by GROMACS version 2020.2^17^ with the PLUMED plugin^18^ used for the targeted MD. The time step of 2 fs was used for equilibration simulations except for the early steps (where it was 1 fs), while targeted and production simulations were performed with a 4-fs time step allowed by repartitioning the mass of heavy atoms into the bonded hydrogen atoms^19^ and the LINCS constraint algorithm^20^. The CHARMM36 force field was used for the protein, lipids, and ions^21^. The topologies for the fluorescent labels were obtained using the SwissParam web-service^22^. They are provided as Supplementary Text 1 and Supplementary Text 2.

### Supplementary figures

#### Fig. S1 Labeling and characterization of the double-cysteine mutant of A_2A_AR.

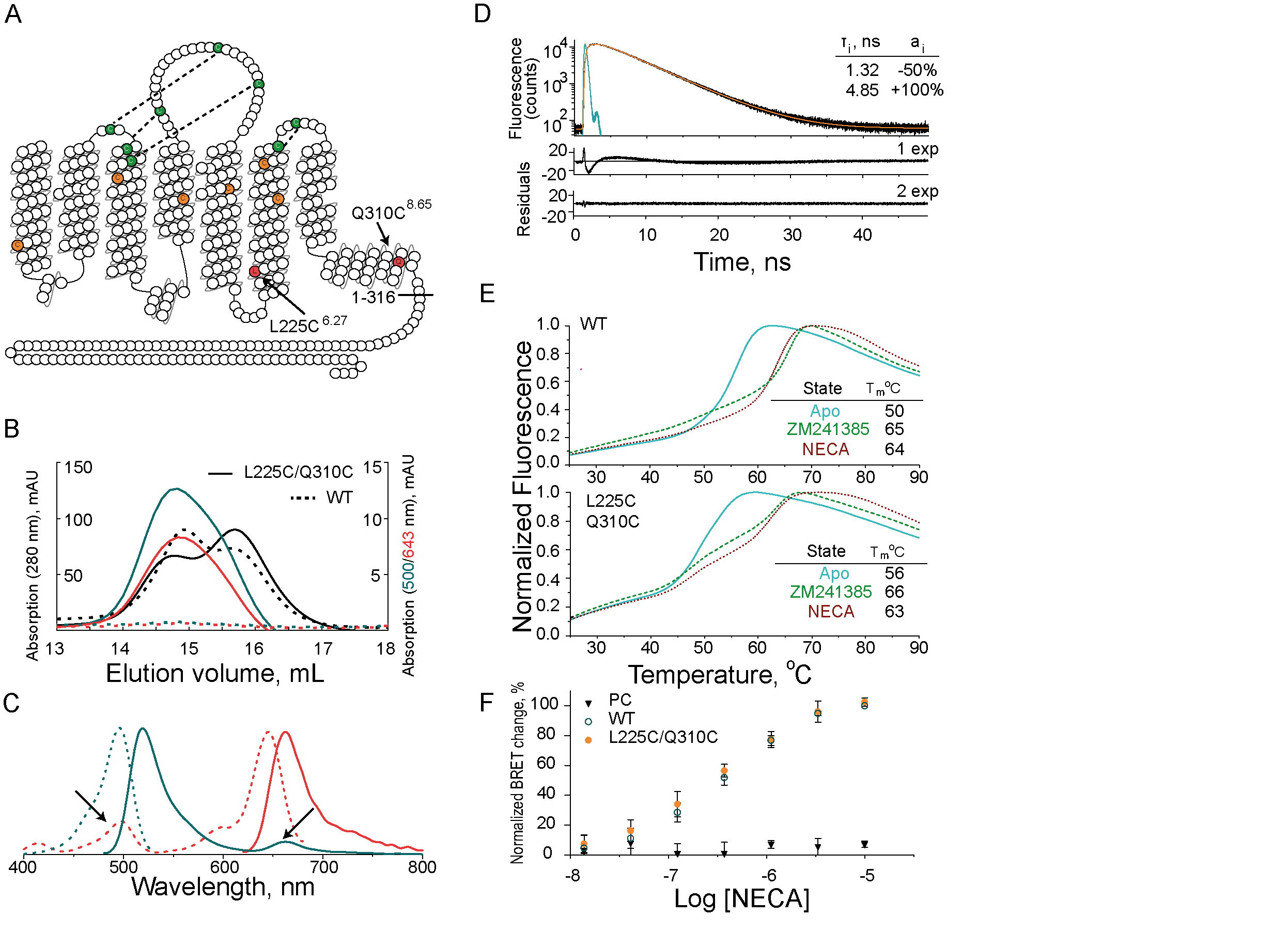

**(A)** A “snake plot” of the amino acid sequence of A_2A_AR (adapted from gpcrdb.org) shows the labelling sites (L225C^6.27^ and Q310C^8.65^) for fluorescent dyes (red circles, black arrows). The native Cys-bridges and unpaired cysteines in A_2A_AR are marked in green and orange, respectively. The C-terminus of A_2A_AR was truncated at residue 316. **(B)** Size-exclusion chromatography of the wild-type (dashed lines) and mutant (solid lines) apo-A_2A_AR in lipid nanodiscs. Dark cyan, red, and black lines correspond to absorption at 500, 643, and 280 nm, respectively. The rightward-shifted peak of the unlabeled protein absorption corresponds to empty nanodiscs. **(C)** Fluorescence excitation (dashed lines) and emission (solid lines) spectra of double-labeled apo-A_2A_AR in lipid nanodiscs indicate FRET (peaks shown with arrows). Donor (dashed, dark cyan) and acceptor (dashed, red) excitation spectra were measured with emission fixed at 550 nm and 700 nm, respectively. Donor (solid, dark cyan) and acceptor (solid, red) emission spectra were measured with excitation fixed at 460 nm and 600 nm, respectively. **(D)** The TCSPC pattern of acceptor emission after pulsed donor excitation shows a rising term associated with FRET in the double-labeled A_2A_AR. Black and orange lines show experimental data and their biexponential fit, respectively. Cyan line shows the IRF (Instrument Response Function) of the setup. Residuals for monoexponential and biexponential fits are shown below the main panel. Parameters of the biexponential fit (fluorescence lifetime and relative amplitudes) are given in the insert table. **(E)** TSA for WT (top frame) and unlabeled double-cysteine mutant (bottom frame) of A_2A_AR bound to ZM241385 (dashed green line), NECA (dashed red line), or apo-A_2A_AR (cyan line). Melting temperature (T_m_) values are given in the insert tables. **(F)** BRET assay shows cAMP signaling induced with agonist NECA in HEK293T cells transfected with a plasmid containing WT (open dark cyan circles) or unlabeled double-cysteine mutant A_2A_AR (closed orange circles) gene. Black triangles show a lack of ligand-response in the vehicle-transfected cells. Data points correspond to the Mean±SD of n = 3 biological replicas. In each experiment, the change in BRET efficiency was normalized to the maximum BRET efficiency change in the WT receptor.

#### Fig. S2. Schematic presentation of the home-built MFD-PIE confocal setup

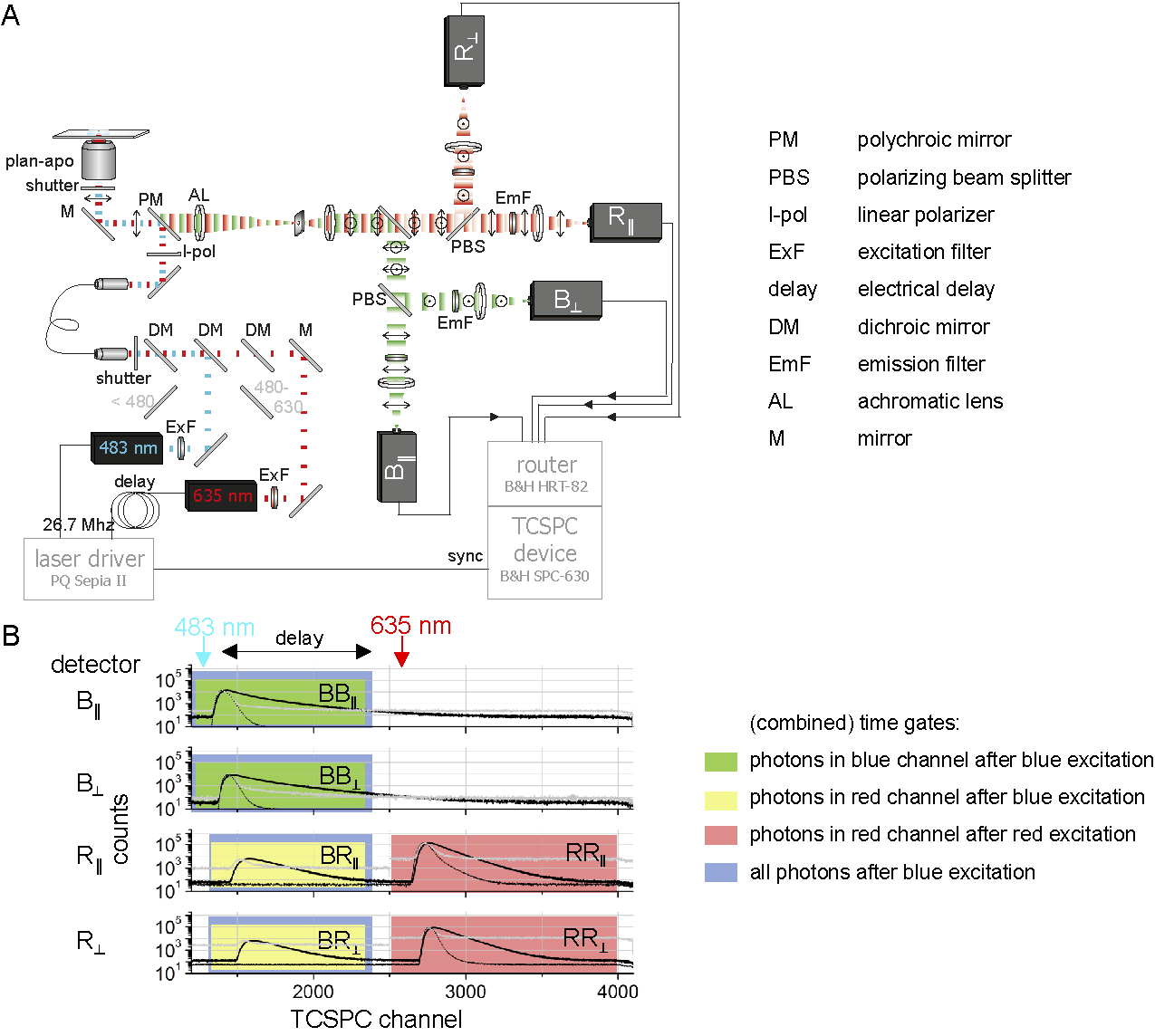

**(A)** Schematic presentation of the home-built MFD-PIE confocal setup^3^. Two pulsed lasers (483 nm and 635 nm) with a ~18 ns lag time, were combined in a single-mode optical fiber, the linear polarization was cleaned up and reflected into the microscope. The transmitted sample emission was focused through a pinhole and spectrally split. Each color (B - blue, R - red) was separately split in two detection channels according to their polarization. Resulting four channels (B_||_, B_⊥_, R_||_, R_⊥_) were detected in TCSPC mode. **(B)** Different time gates (‘PIE channels’) in the previously described detection channels depending on the blue or red excitation line. Data (thick black line), IRF (thin black line), background recording (gray) are illustrated, along with the nomenclature of the different detectors and time gates. Since the 635 nm laser is delayed, first photons are detected on the BB_||_/BB_⊥_ (donor direct excitation) and BR_║_/BR_⊥_ (FRET-channel) time gates. After the delay, 635 nm excitation triggers photons in the RR_║_ and RR_⊥_ time gates (direct excitation of an acceptor).

#### Fig. S3. Stoichiometry and ALEX-2CDE filters isolate fluorescence bursts corresponding to the double-labeled receptor subpopulation in single-molecule experiments.

| A.   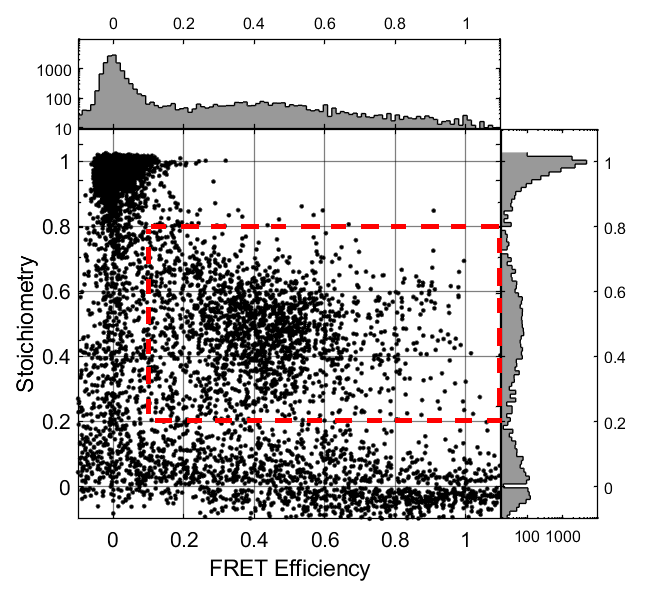 | B.  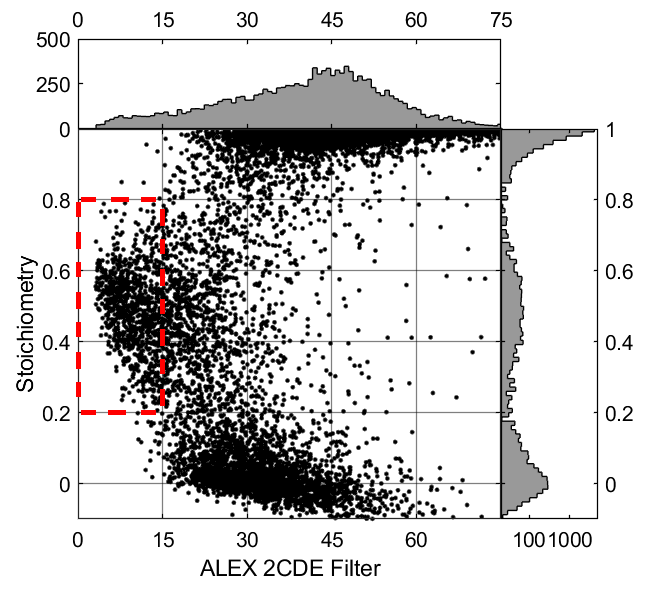 |
| --- | --- |

**(A)** Scatter plot of stoichiometry S against FRET efficiency – the donor-only molecules are centered around the top-left corner, the acceptor-only molecules cluster in the bottom of the plot space. The double-labeled molecules in the red rectangle were selected for the further analysis. **(B)** Scatter plot of stoichiometry *S* against ALEX-2CDE filter value. The donor-only and acceptor-only molecules tend to the top-right and bottom-right parts of the plot, respectively. The double-labeled molecules in the red rectangle were selected for further analysis.

#### Fig. S4. Microtime patterns (top) and filter functions (bottom) used for fFCS.

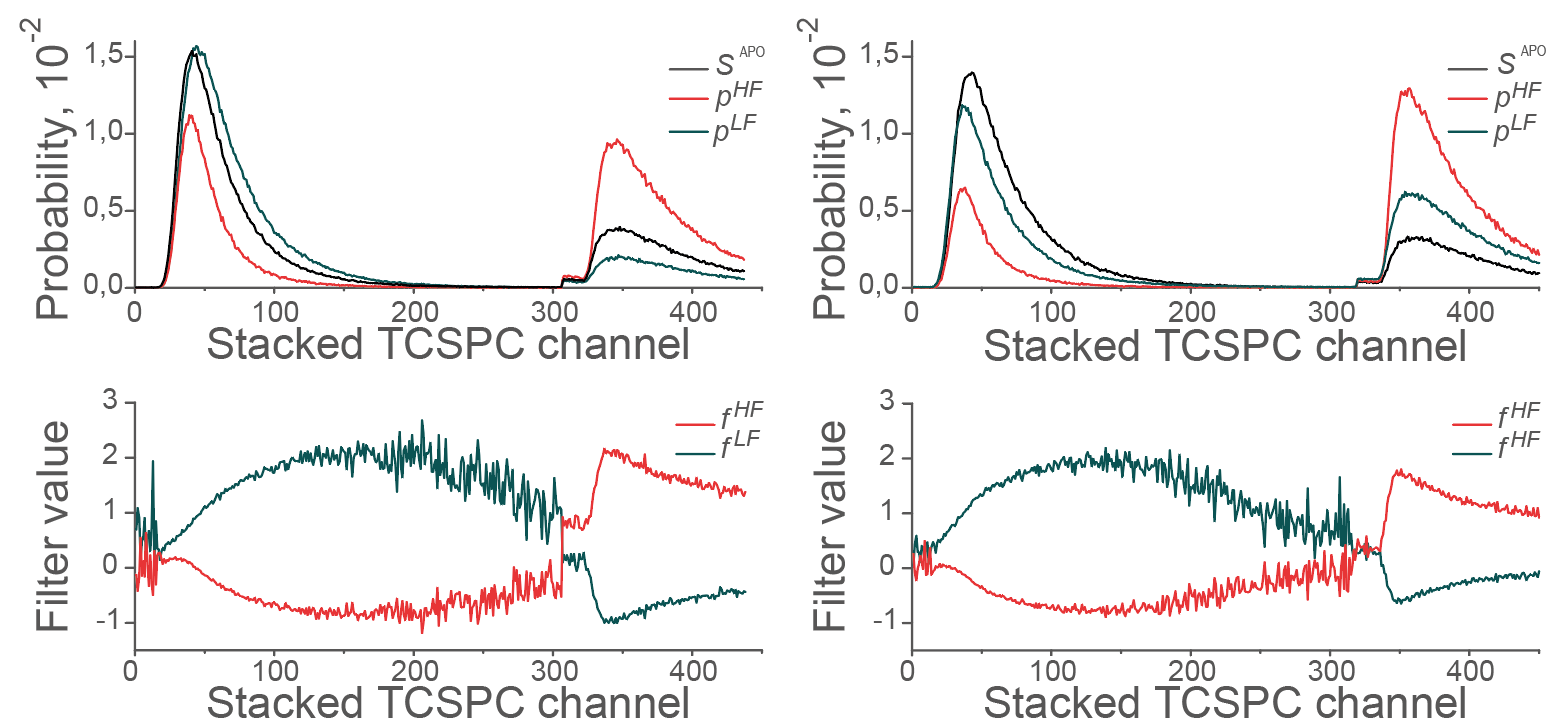

The left and right panels correspond to the channels of parallel and perpendicular fluorescence polarization, respectively. *S^APO^* (black lines) is the microtime-distribution of photons accumulated from the double-labeled apo-A_2A_AR molecules. *p^LF^* (red, top pannel) and *p^HF^* (dark green, top panel) show TCSPC-distributions accumulated from bursts with low (E<0.3) and high (E>0.7) burst-wise FRET efficiency (all ligand-bound and apo-conditions merged). *f ^LF^* and *f ^HF^* show filter functions used for the ‘low-FRET’ and ‘high-FRET’ channels in fFCS with the apo-A_2A_AR.

#### Fig. S5 PDA histograms for interconverting medium-FRET and high-FRET states.

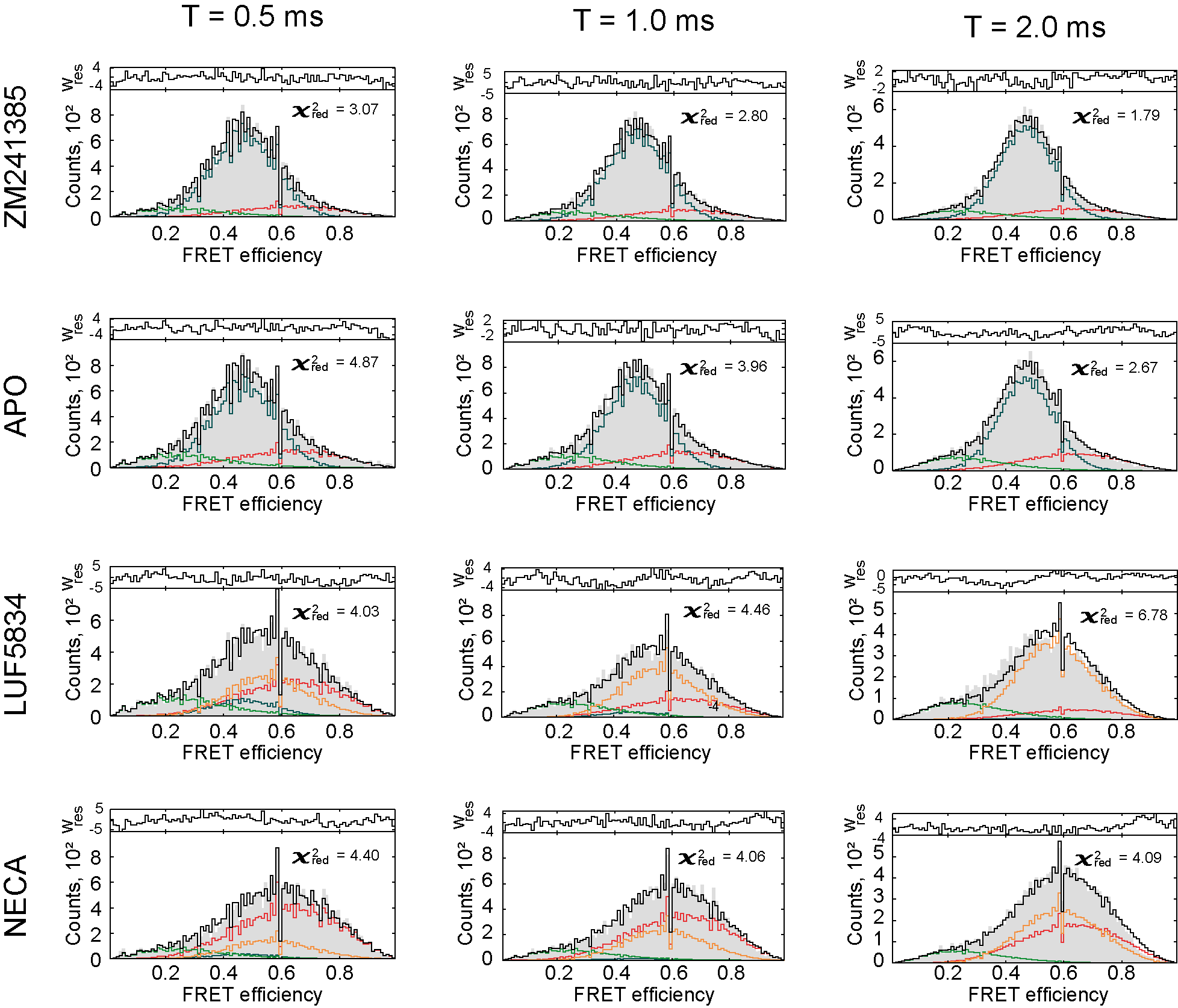

PDA histograms were fit with a sum of three states, allowing the medium-FRET (dark-green) and high-FRET (red) states to interconvert. The longer exchange time derived from fFCS (τ_2_) was used for LUF5834 and NECA. The infinitesimally long exchange times (~100 ms) were used for ZM241385 and apo-condition. Columns show PDA distributions for three different time-bin lengths, rows correspond to A_2A_AR in different ligand-bound or apo-conditions. The experimental distributions are shown as grey bars. The resulting fit (black line) is a sum of distributions simulated for molecules that stay in the low-FRET (light green line), medium-FRET (dark cyan line), or high-FRET (red line) state during the entire simulated time-bin, and the distribution for molecules that sample both medium-FRET and high-FRET states within a time-bin (orange line). The fitting residuals are given on the top of each panel. For the global fit, χ^2^_red_ = 3.6.

#### Fig. S6 PDA histograms for ‘static’ states.

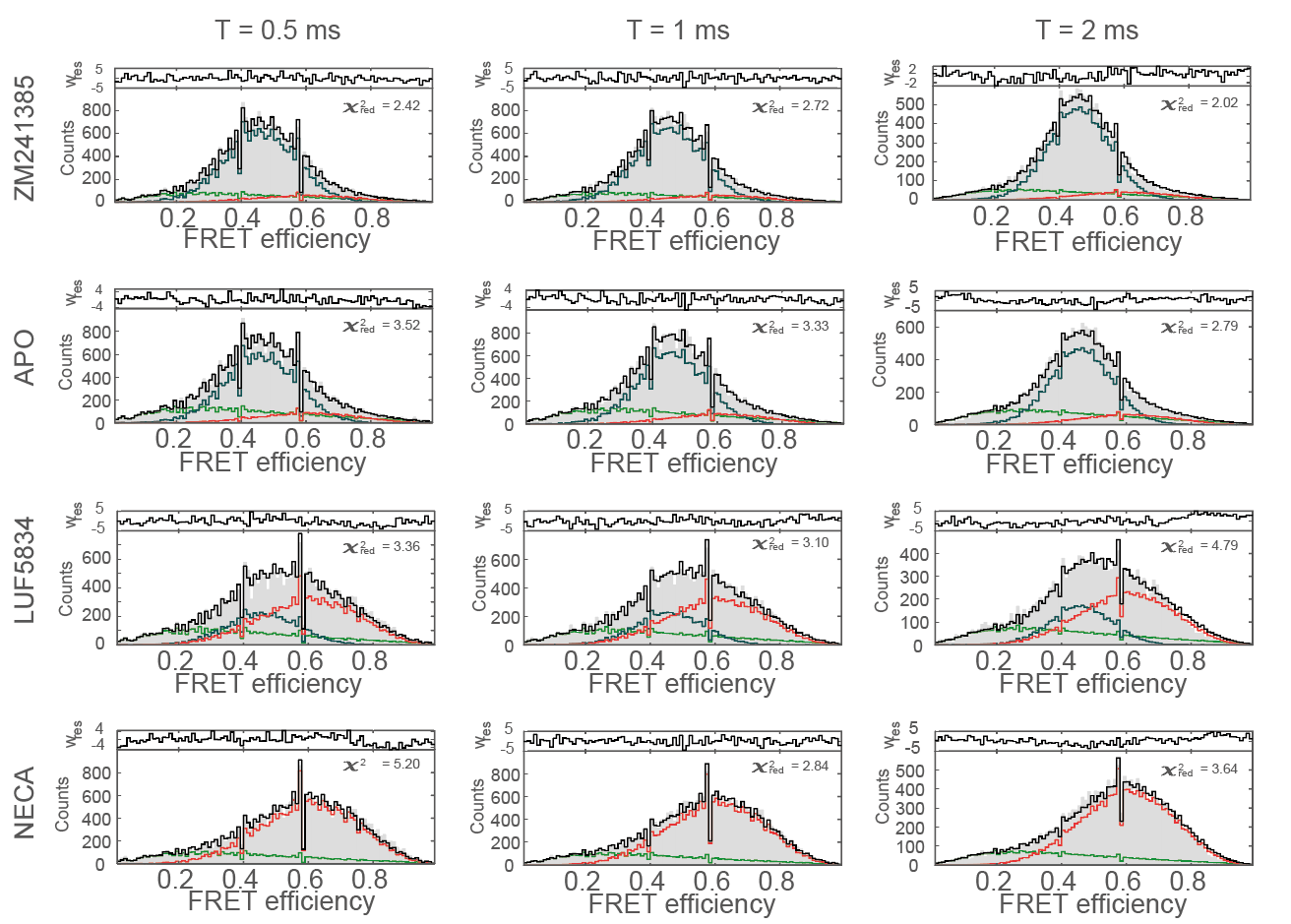

PDA histograms were fit with a sum of three non-interconverting (‘static’) states. Columns show PDA distributions for three different time-bin lengths; rows correspond to A_2A_AR in different ligand-bound or apo-conditions. The fitting curve (black line) is shown on top of the experimental distributions (grey bars). The fitting residuals are given on the top of each panel. Simulated distributions for individual states are shown in light green (low-FRET), dark cyan (medium-FRET), and red (high-FRET) lines. For the global fit, χ ^2^_red_ = 3.0.

#### Fig. S7. Unconstrained dynamic PDA

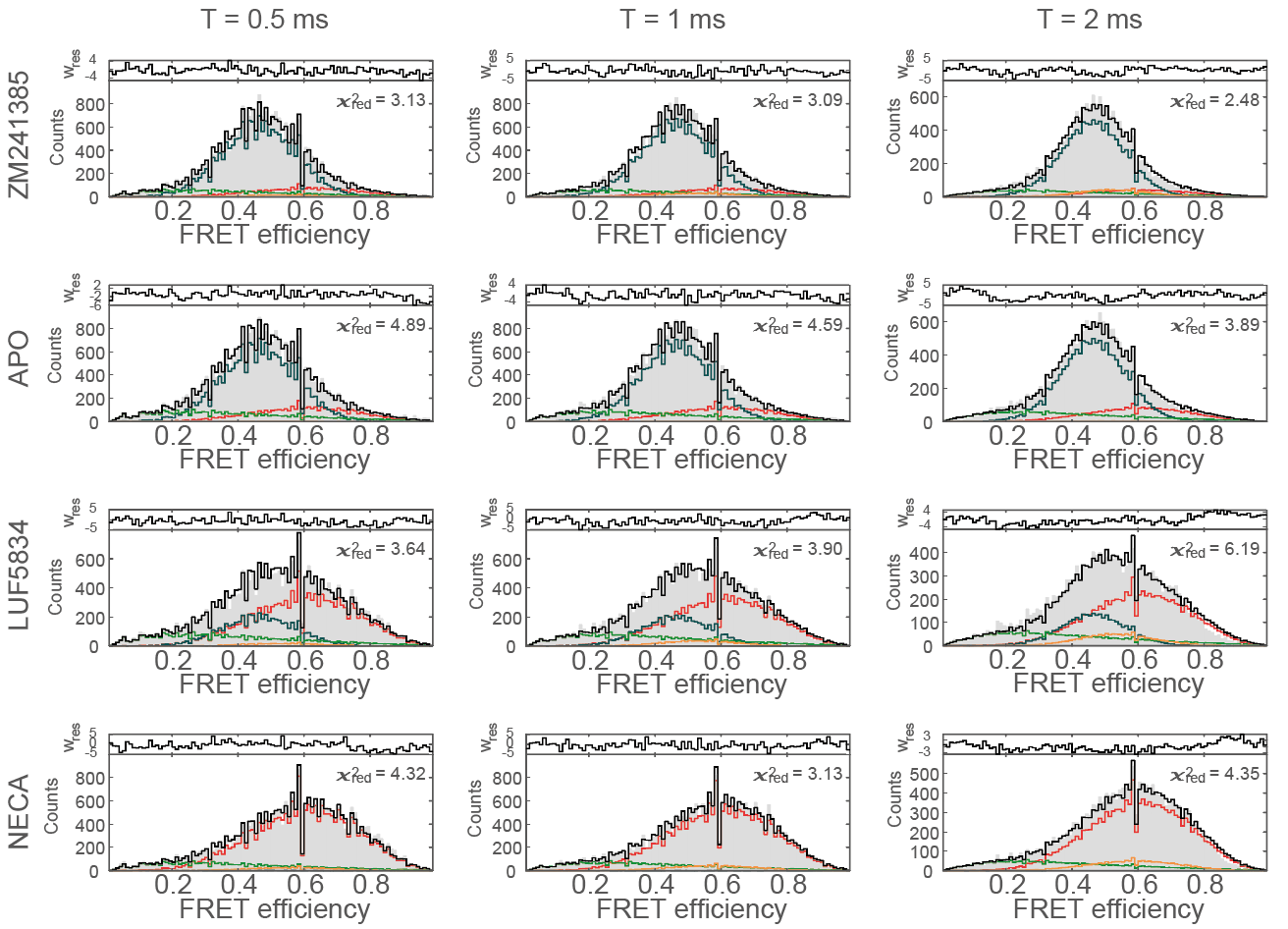

PDA histograms were fit with a sum of three states, allowing the medium-FRET (dark-green) and high-FRET (red) states to interconvert. The exchange time was set as a free fit parameter. Columns show PDA distributions for three different time-bin lengths, rows correspond to A_2A_AR in different ligand-bound or apo-conditions. The experimental distributions are shown as grey bars. The resulting fit (black line) is a sum of distributions simulated for molecules that stay in the low-FRET (light green line), medium-FRET (dark cyan line), or high-FRET (red line) state during the entire simulated time-bin, and the distribution for molecules that sample both medium-FRET and high-FRET states within a time-bin (orange line). The fitting residuals are given on the top of each panel. For the global fit, χ ^2^_red_ = 3.6.

#### Fig. S8. Targeted-MD simulations suggest that the dye attached to TM6 can enter the G-protein-binding cavity upon A_2A_AR activation.

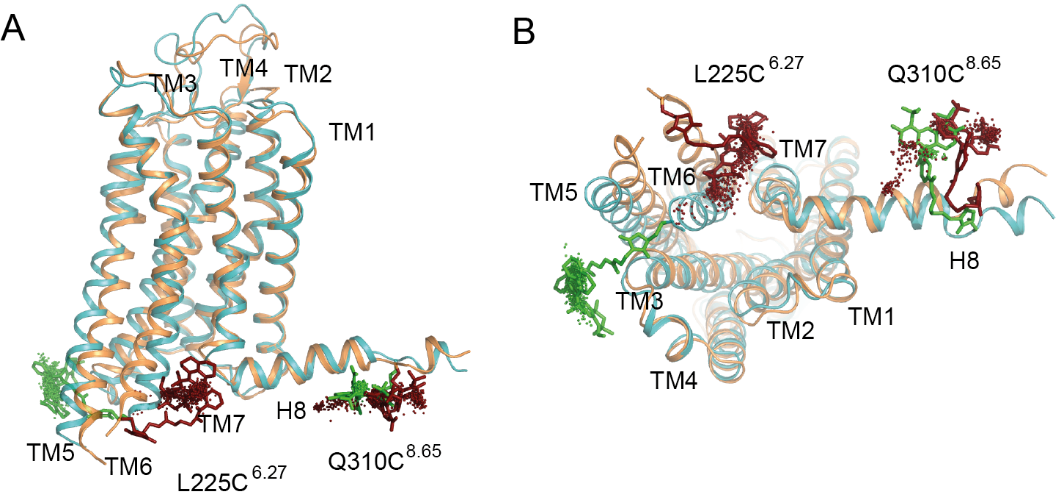

The inactive (PDB: 3PWH) and fully active (PDB: 5G53) structures of A_2A_AR after equilibration are shown in cyan and orange, respectively. Representative positions of the dyes are shown with either green or red sticks for inactive and active structures, respectively. Dots indicate the geometrical centers of the fluorophores sampled in 1-μs-long MD trajectories with 1-ns time-steps for each A_2A_AR state. Here, Atto643 is attached to L225C^6.27^, Alexa 488 is attached to Q310C^8.65^, the simulation of the alternative labeling variant showed similar results.

#### Table S1. Labeling efficiencies

| Mutations | Alexa 488, % | Atto 643, % |
| --- | --- | --- |
| Q310C^8.65^/L225C^6.27^ | 26 | 8 |
| WT | 1.1 | 0.4 |

Labeling efficiencies of the wild type and double-cysteine mutant of A_2A_AR with Alexa 488 and Atto 643 dyes based on the corresponding absorption, assuming one binding site for each dye on a single protein.

#### Table S2. Mean burst-wise parameters

|  | ZM241385 | APO | LUF5834 | NECA |
| --- | --- | --- | --- | --- |
| r_D_ | 0.173±0.001 | 0.173±0.002 | 0.191±0.004 | 0.190±0.004 |
| t_D_ | 3.77±0.02 | 3.80±0.02 | 3.75±0.01 | 3.74±0.01 |
| r_D(A)_ | 0.195±0.002 | 0.202±0.002 | 0.211±0.002 | 0.213±0.006 |
| t_D(A)_ | 2.50±0.06 | 2.50±0.04 | 2.42±0.11 | 2.20±0.02 |
| E | 0.45±0.01 | 0.45±0.01 | 0.50±0.02 | 0.55±0.01 |
| r_A(D)_ | 0.261±0.002 | 0.253±0.004 | 0.260±0.001 | 0.261±0.002 |
| t_A(D)_ | 4.57±0.02 | 4.55±0.08 | 4.56±0.02 | 4.53±0.01 |
| FRET-2CDE | 13.8±0.2 | 14.0±0.2 | 16.1±0.5 | 15.8±0.3 |
| r_A_ | 0.261±0.003 | 0.252±0.004 | 0.265±0.002 | 0.263±0.001 |
| t_A_ | 4.51±0.04 | 4.47±0.11 | 4.45±0.05 | 4.43±0.02 |

Mean burst-wise parameters are given with S.D. between the mean parameter values of n  =  3 independent experiments. Donor and acceptor fluorescence lifetime (t_D,_ t_A_) and anisotropy (r_D_, r_A_) were determined for single-labeled molecules. For double-labeled molecules, donor fluorescence lifetime (t_D(A)_) and anisotropy (r_D(A)_), FRET efficiency (E), acceptor fluorescence lifetime (t_A(D)_) and acceptor anisotropy (r_A(D_) were determined.

#### Table S3. Fitting parameters for fFCS cross-correlation curves.

(A)

|  | T_D_, ms | A_1_ | A_2_ | τ_1_, μs | τ_2_, μs |
| --- | --- | --- | --- | --- | --- |
| ZM241385 | 1.22±0.05 | 0.6±0.2 | 0.10±0.02 | 2±1 | 80±40 |
| APO | 1.1±0.2 | 0.32±0.06 | 0.13±0.06 | 9±3 | 300±200 |
| LUF5834 | 2.1±0.2 | 0.6±0.1 | 0.35±0.05 | 7±2 | 100±30 |
| NECA | 1.5±0.2 | 0.5±0.1 | 0.35±0.05 | 10±3 | 150±50 |

(B)

|  | T_D_, ms | A_1_ | A_2_ | T_1_, μs | T_2_, μs |
| --- | --- | --- | --- | --- | --- |
| ZM241385 | 1.38±0.04 | 0.5±0.2 | - | 3±1 | - |
| APO | 1.38±0.04 | 0.38±0.06 | - | 14±4 | - |
| LUF5834 | 1.38±0.04 | 0.54±0.05 | 0.36±0.04 | 18±4 | 390±80 |
| NECA | 1.38±0.04 | 0.38±0.08 | 0.37±0.04 | 16±6 | 390±80 |

**(A)** The fitting was performed without any constrains; two dynamic terms were found for the apo and agonist-bound receptor. **(B)** In the final model, the second dynamics term (A_2_) was set non-zero only for the agonist-bound A_2A_AR; T_D_ and T_2_ were optimized globally across the apo and three ligand-bound conditions. The half-widths of 95% confidence intervals are given as fitting errors.

#### Table S4 Fitting PDA parameters for a model with one long-living and two interconverting states (the exchange time was derived from the fFCS).

| (A) |  |  |  |  |
| --- | --- | --- | --- | --- |
|  | R, Å | σ, Å |  |  |
| LF | 57.9 | 6.0 |  |  |
| MF | 50.0 | 2.1 |  |  |
| HF | 45.1 | 4.9 |  |  |
| (B) |  |  |  |  |
|  | T_ex_, ms | LF, % | MF, % | HF, % |
| ZM241385 | > 2 | 11±3 | 74±3 | 15±1 |
| APO | > 2 | 15±5 | 64±6 | 20±3 |
| LUF5834 | 0.39 | 20±1 | 25±8 | 55±8 |
| NECA | 0.39 | 12±4 | 11±1 | 77±3 |

**(A)** The means and standard deviations of the Gaussian distance distributions for individual states (low-FRET - LF, medium FRET - MF, high FRET -HF). **(B)** Populations (mean ± SD of three independent experiments) of the three states as well as their exchange times (columns) for the ligand-bound or apo-A_2A_AR (rows).

#### Table S5. Fitting PDA parameters for a model with three non-interconverting (static) states.

| (A) |  |  |  |
| --- | --- | --- | --- |
|  | R, Å | σ, Å |  |
| LF | 53.4 | 9.1 |  |
| MF | 50.3 | 2.1 |  |
| HF | 45.8 | 4.1 |  |
| (B) |  |  |  |
|  | LF, % | MF, % | HF, % |
| ZM241385 | 19 | 72 | 9 |
| APO | 27 | 61 | 12 |
| LUF5834 | 25 | 25 | 50 |
| NECA | 22 | 0 | 78 |

**(A)** The means and standard deviations of the Gaussian distance distributions for individual states (low-FRET - LF, medium FRET - MF, high FRET -HF). **(B)** Populations of the three states (columns) for the ligand-bound or apo-A_2A_AR (rows).

#### Table S6. Fitting PDA parameters for a model with one long-living and two interconverting states with a free exchange time.

| (A) |  |  |  |  |
| --- | --- | --- | --- | --- |
|  | R, A | σ, A |  |  |
| LF | 54.3 | 9.2 |  |  |
| MF | 50.2 | 2.3 |  |  |
| HF | 45.7 | 4.4 |  |  |
| (B) |  |  |  |  |
|  | T_ex_, ms | LF, % | MF, % | HF, % |
| ZM241385 | 5.5 | 15 | 72 | 13 |
| APO | 100 | 20 | 63 | 17 |
| LUF5834 | 7.2 | 22 | 23 | 55 |
| NECA | 0.26 | 17 | 1 | 82 |

Fitting PDA parameters for a model with one long-living and two interconverting states with a free exchange time. **(A)** The means and standard deviations of the Gaussian distance distributions for individual states (low-FRET - LF, medium FRET - MF, high FRET -HF). **(B)** Populations of the three states as well as the optimal exchange times (columns) for the ligand-bound or apo-A_2A_AR (rows).

**Supplementary text 1.** Topology for Atto647N-maleimide. (uploaded separately)

**Supplementary text 2.** Topology for Alexa488-C5-maleimide. (uploaded separately)
