## Supplementary text 1. Topology for Atto647N-maleimide. for "Sub-millisecond conformational dynamics of the A_2A_ adenosine receptor revealed by single-molecule FRET"

**GROMACS rtp file**

[ ATT ]

[ atoms ]

C1	CG2DC1	-0.160	1

C2	CG2DC1	0.035	2

C3	CG2DC2	0.556	3

N4	NG2P1	-0.400	4

C5	CG324	0.232	5

C6	CG321	-0.162	6

C7	CG321	-0.197	7

C8	CG324	0.232	8

C9	CG321	-0.162	9

C10	CG321	-0.205	10

C11	CG2DC1	0.016	11

C12	CG2DC1	0.000	12

C13	CG2DC2	-0.008	13

C14	CG2DC2	0.009	14

C15	CG2R61	0.002	15

C16	CG2R61	-0.003	16

C17	CG301	0.001	17

C18	CG331	-0.271	18

C19	CG331	-0.271	19

C20	CG2R61	-0.111	20

C21	CG2R61	0.105	21

C22	CG2R61	-0.001	22

C23	CG2R61	-0.113	23

C24	CG311	-0.082	24

C25	CG321	-0.186	25

C26	CG301	0.056	26

N27	NG301	-0.338	27

C28	CG321	0.004	28

C29	CG331	-0.273	29

C30	CG331	-0.273	30

C31	CG331	-0.273	31

C32	CG331	-0.272	32

C33	CG2R61	0.032	33

C34	CG2R61	-0.131	34

C35	CG2R61	-0.067	35

C36	CG2R61	-0.100	36

C37	CG2R61	-0.115	37

C38	CG2R61	-0.100	38

C39	CG2O1	0.424	39

O40	OG2D1	-0.449	40

N41	NG2S0	-0.438	41

C42	CG331	-0.063	42

C43	CG321	0.094	43

C44	CG321	-0.167	44

C45	CG321	-0.227	45

C46	CG2O1	0.509	46

O47	OG2D1	-0.511	47

N48	NG2S1	-0.494	48

C49	CG321	0.020	49

C50	CG321	0.039	50

N51	NG2R53	-0.171	51

C52	CG2R53	0.312	52

O53	OG2D1	-0.487	53

C54	CG3C52	-0.028	54

C55	CG3C51	0.160	55

C56	CG2R53	0.314	56

O57	OG2D1	-0.487	57

S58	SG311	-0.156	58

C59	CT2	-0.151	59

CA	CT1	0.383	60

C	C	0.448	61

O	O	-0.860	62

H63	HGA4	0.150	63

H64	HGA2	0.090	64

H65	HGA2	0.090	65

H66	HGA2	0.090	66

H67	HGA2	0.090	67

H68	HGA2	0.090	68

H69	HGA2	0.090	69

H70	HGA2	0.090	70

H71	HGA2	0.090	71

H72	HGA2	0.090	72

H73	HGA2	0.090	73

H74	HGA2	0.090	74

H75	HGA2	0.090	75

H76	HGA3	0.090	76

H77	HGA3	0.090	77

H78	HGA3	0.090	78

H79	HGA3	0.090	79

H80	HGA3	0.090	80

H81	HGA3	0.090	81

H82	HGR61	0.115	82

H83	HGR61	0.115	83

H84	HGA1	0.090	84

H85	HGA2	0.090	85

H86	HGA2	0.090	86

H87	HGA2	0.090	87

H88	HGA2	0.090	88

H89	HGA3	0.090	89

H90	HGA3	0.090	90

H91	HGA3	0.090	91

H92	HGA3	0.090	92

H93	HGA3	0.090	93

H94	HGA3	0.090	94

H95	HGA3	0.090	95

H96	HGA3	0.090	96

H97	HGA3	0.090	97

H98	HGA3	0.090	98

H99	HGA3	0.090	99

H100	HGA3	0.090	100

H101	HGR61	0.115	101

H102	HGR61	0.115	102

H103	HGR61	0.115	103

H104	HGR61	0.115	104

H105	HGA3	0.090	105

H106	HGA3	0.090	106

H107	HGA3	0.090	107

H108	HGA2	0.090	108

H109	HGA2	0.090	109

H110	HGA2	0.090	110

H111	HGA2	0.090	111

H112	HGA2	0.090	112

H113	HGA2	0.090	113

H114	HGP1	0.308	114

H115	HGA2	0.090	115

H116	HGA2	0.090	116

H117	HGA2	0.090	117

H118	HGA2	0.090	118

H119	HGA2	0.090	119

H120	HGA2	0.090	120

H121	HGA1	0.090	121

H122	HA2	0.090	122

H123	HA2	0.090	123

HA	HB1	0.090	124

N	NH1	-0.358	125

HN	H	0.330	126

[ bonds ]

C1 C13

C1 C2

C1 H63

C2 C7

C2 C3

C3 C11

C3 N4

N4 C5

N4 C8

C5 C6

C5 H64

C5 H65

C6 C7

C6 H66

C6 H67

C7 H68

C7 H69

C8 C9

C8 H70

C8 H71

C9 C10

C9 H72

C9 H73

C10 C11

C10 H74

C10 H75

C11 C12

C12 C17

C12 C13

C13 C14

C14 C15

C14 C33

C15 C23

C15 C16

C16 C17

C16 C20

C17 C18

C17 C19

C18 H76

C18 H77

C18 H78

C19 H79

C19 H80

C19 H81

C20 C21

C20 H82

C21 N27

C21 C22

C22 C23

C22 C24

C23 H83

C24 C25

C24 C32

C24 H84

C25 C26

C25 H85

C25 H86

C26 N27

C26 C30

C26 C31

N27 C28

C28 C29

C28 H87

C28 H88

C29 H89

C29 H90

C29 H91

C30 H92

C30 H93

C30 H94

C31 H95

C31 H96

C31 H97

C32 H98

C32 H99

C32 H100

C33 C38

C33 C34

C34 C35

C34 C39

C35 C36

C35 H101

C36 C37

C36 H102

C37 C38

C37 H103

C38 H104

C39 O40

C39 N41

N41 C42

N41 C43

C42 H105

C42 H106

C42 H107

C43 C44

C43 H108

C43 H109

C44 C45

C44 H110

C44 H111

C45 C46

C45 H112

C45 H113

C46 O47

C46 N48

N48 C49

N48 H114

C49 C50

C49 H115

C49 H116

C50 N51

C50 H117

C50 H118

N51 C56

N51 C52

C52 O53

C52 C54

C54 C55

C54 H119

C54 H120

C55 C56

C55 S58

C55 H121

C56 O57

S58 C59

C59 CA

C59 H122

C59 H123

CA C

CA HA

CA N

C	O

N HN

C +N

**GROMACS prm file**

[ bondtypes ]

; i	j func	b0	kb

CG2DC1	CG2DC1	1	0.13400000	368192.00

CG2DC1	CG2DC2	1	0.14500000	251040.00

CG2DC1	CG301	1	0.15020000	305432.00

CG2DC1	CG321	1	0.15020000	305432.00

CG2DC1	HGA4	1	0.11000000	301666.40

CG2DC2	CG2DC2	1	0.13400000	368192.00

CG2DC2	CG2R61	1	0.14500000	305432.00

CG2DC2	NG2P1	1	0.12830000	393296.00

CG2O1	CG2R61	1	0.14750000	251040.00

CG2O1	CG321	1	0.14900000	209200.00

CG2O1	NG2S0	1	0.13500000	359824.00

CG2O1	NG2S1	1	0.13450000	309616.00

CG2O1	OG2D1	1	0.12300000	518816.00

CG2O3	CG314	1	0.15220000	167360.00

CG2O3	OG2D2	1	0.12600000	439320.00

CG2R53	CG3C51	1	0.15300000	251040.00

CG2R53	CG3C52	1	0.15300000	251040.00

CG2R53	NG2R53	1	0.13800000	384928.00

CG2R53	OG2D1	1	0.12350000	476976.00

CG2R61	CG2R61	1	0.13750000	255224.00

CG2R61	CG301	1	0.14900000	192464.00

CG2R61	CG311	1	0.14900000	192464.00

CG2R61	NG301	1	0.14100000	271960.00

CG2R61	HGR61	1	0.10800000	284512.00

CG301	CG321	1	0.15380000	186188.00

CG301	CG331	1	0.15380000	186188.00

CG301	NG301	1	0.14500000	167360.00

CG311	CG321	1	0.15380000	186188.00

CG311	CG331	1	0.15380000	186188.00

CG311	HGA1	1	0.11110000	258571.20

CG314	CG321	1	0.15380000	186188.00

CG314	NG3P3	1	0.14800000	167360.00

CG314	HGA1	1	0.11110000	258571.20

CG321	CG321	1	0.15300000	186188.00

CG321	CG324	1	0.15300000	186188.00

CG321	CG331	1	0.15280000	186188.00

CG321	NG2R53	1	0.14300000	267776.00

CG321	NG2S1	1	0.14300000	267776.00

CG321	NG301	1	0.14500000	167360.00

CG321	SG311	1	0.18180000	165686.40

CG321	HGA2	1	0.11110000	258571.20

CG324	NG2P1	1	0.14530000	251040.00

CG324	HGA2	1	0.11000000	238069.60

CG331	NG2S0	1	0.14340000	263592.00

CG331	HGA3	1	0.11110000	269449.60

CG3C51	CG3C52	1	0.15180000	163176.00

CG3C51	SG311	1	0.18180000	165686.40

CG3C51	HGA1	1	0.11000000	256897.60

CG3C52	HGA2	1	0.11000000	256897.60

NG2S1	HGP1	1	0.09970000	368192.00

NG3P3	HGP2	1	0.10400000	337230.40

[ angletypes ]

; i	j	k func	theta0	ktheta	ub0	kub

CG2DC1	CG2DC1	CG2DC2	5	123.000000	401.664000	0.00000000	0.00

CG2DC1	CG2DC1	CG301	5	123.500000	401.664000	0.00000000	0.00

CG2DC1	CG2DC1	CG321	5	123.500000	401.664000	0.00000000	0.00

CG2DC1	CG2DC1	HGA4	5	119.000000	351.456000	0.00000000	0.00

CG2DC2	CG2DC1	CG301	5	123.500000	401.664000	0.00000000	0.00

CG2DC2	CG2DC1	CG321	5	113.000000	401.664000	0.00000000	0.00

CG2DC2	CG2DC1	HGA4	5	118.000000	351.456000	0.00000000	0.00

CG2DC1	CG2DC2	CG2DC1	5	113.000000	401.664000	0.00000000	0.00

CG2DC1	CG2DC2	CG2DC2	5	123.000000	401.664000	0.00000000	0.00

CG2DC1	CG2DC2	NG2P1	5	125.600000	334.720000	0.00000000	0.00

CG2DC2	CG2DC2	CG2R61	5	122.000000	242.672000	0.00000000	0.00

CG2R61	CG2DC2	CG2R61	5	113.000000	401.664000	0.00000000	0.00

CG2R61	CG2O1	NG2S0	5	116.500000	669.440000	0.00000000	0.00

CG2R61	CG2O1	OG2D1	5	121.000000	251.040000	0.00000000	0.00

CG321	CG2O1	NG2S1	5	116.500000	669.440000	0.00000000	0.00

CG321	CG2O1	OG2D1	5	121.000000	669.440000	0.00000000	0.00

NG2S0	CG2O1	OG2D1	5	124.000000	669.440000	0.00000000	0.00

NG2S1	CG2O1	OG2D1	5	122.500000	669.440000	0.00000000	0.00

CG314	CG2O3	OG2D2	5	116.000000	334.720000	0.23530000	41840.00

OG2D2	CG2O3	OG2D2	5	128.000000	836.800000	0.22587000	58576.00

CG3C51	CG2R53	NG2R53	5	105.500000 1004.160000	0.00000000	0.00

CG3C51	CG2R53	OG2D1	5	126.700000	543.920000	0.00000000	0.00

CG3C52	CG2R53	NG2R53	5	105.500000 1004.160000	0.00000000	0.00

CG3C52	CG2R53	OG2D1	5	126.700000	543.920000	0.00000000	0.00

NG2R53	CG2R53	OG2D1	5	127.800000	543.920000	0.00000000	0.00

CG2DC2	CG2R61	CG2R61	5	120.000000	301.248000	0.00000000	0.00

CG2O1	CG2R61	CG2R61	5	119.000000	376.560000	0.00000000	0.00

CG2R61	CG2R61	CG2R61	5	120.000000	334.720000	0.24162000	29288.00

CG2R61	CG2R61	CG301	5	120.000000	383.254400	0.00000000	0.00

CG2R61	CG2R61	CG311	5	120.000000	383.254400	0.00000000	0.00

CG2R61	CG2R61	NG301	5	120.000000	351.456000	0.00000000	0.00

CG2R61	CG2R61	HGR61	5	120.000000	251.040000	0.21525000	18409.60

CG2DC1	CG301	CG2R61	5	107.500000	433.462400	0.00000000	0.00

CG2DC1	CG301	CG331	5	112.200000	267.776000	0.00000000	0.00

CG2R61	CG301	CG331	5	107.500000	433.462400	0.00000000	0.00

CG321	CG301	CG331	5	113.500000	488.272800	0.25610000	9338.69

CG321	CG301	NG301	5	107.000000	476.976000	0.00000000	0.00

CG331	CG301	CG331	5	113.500000	488.272800	0.25610000	9338.69

CG331	CG301	NG301	5	112.200000	365.681600	0.00000000	0.00

CG2R61	CG311	CG321	5	107.500000	433.462400	0.00000000	0.00

CG2R61	CG311	CG331	5	107.500000	433.462400	0.00000000	0.00

CG2R61	CG311	HGA1	5	111.000000	359.824000	0.00000000	0.00

CG321	CG311	CG331	5	114.000000	446.432800	0.25610000	6694.40

CG321	CG311	HGA1	5	110.100000	288.696000	0.21790000	18853.10

CG331	CG311	HGA1	5	110.100000	288.696000	0.21790000	18853.10

CG2O3	CG314	CG321	5	108.000000	435.136000	0.00000000	0.00

CG2O3	CG314	NG3P3	5	110.000000	365.681600	0.00000000	0.00

CG2O3	CG314	HGA1	5	109.500000	418.400000	0.00000000	0.00

CG321	CG314	NG3P3	5	110.000000	566.513600	0.00000000	0.00

CG321	CG314	HGA1	5	110.100000	288.696000	0.21790000	18853.10

NG3P3	CG314	HGA1	5	107.500000	430.952000	0.00000000	0.00

CG2DC1	CG321	CG321	5	112.200000	267.776000	0.00000000	0.00

CG2DC1	CG321	HGA2	5	111.500000	376.560000	0.00000000	0.00

CG2O1	CG321	CG321	5	108.000000	435.136000	0.00000000	0.00

CG2O1	CG321	HGA2	5	109.500000	276.144000	0.21630000	25104.00

CG301	CG321	CG311	5	113.500000	488.272800	0.25610000	9338.69

CG301	CG321	HGA2	5	110.100000	221.752000	0.21790000	18853.10

CG311	CG321	HGA2	5	110.100000	279.742240	0.21790000	18853.10

CG314	CG321	SG311	5	112.500000	485.344000	0.00000000	0.00

CG314	CG321	HGA2	5	110.100000	279.742240	0.21790000	18853.10

CG321	CG321	CG321	5	113.600000	488.272800	0.25610000	9338.69

CG321	CG321	CG324	5	110.500000	488.272800	0.25610000	9338.69

CG321	CG321	NG2R53	5	113.500000	585.760000	0.00000000	0.00

CG321	CG321	NG2S1	5	113.500000	585.760000	0.00000000	0.00

CG321	CG321	HGA2	5	110.100000	221.752000	0.21790000	18853.10

CG324	CG321	HGA2	5	110.100000	221.752000	0.21790000	18853.10

CG331	CG321	NG301	5	112.200000	365.681600	0.00000000	0.00

CG331	CG321	HGA2	5	110.100000	289.532800	0.21790000	18853.10

NG2R53	CG321	HGA2	5	109.500000	430.952000	0.00000000	0.00

NG2S1	CG321	HGA2	5	109.500000	430.952000	0.00000000	0.00

NG301	CG321	HGA2	5	109.000000	271.123200	0.00000000	0.00

SG311	CG321	HGA2	5	111.300000	385.764800	0.00000000	0.00

HGA2	CG321	HGA2	5	109.000000	297.064000	0.18020000	4518.72

CG321	CG324	NG2P1	5	110.000000	566.513600	0.00000000	0.00

CG321	CG324	HGA2	5	111.800000	221.752000	0.21790000	18853.10

NG2P1	CG324	HGA2	5	110.100000	351.456000	0.00000000	0.00

HGA2	CG324	HGA2	5	109.000000	297.064000	0.18020000	4518.72

CG301	CG331	HGA3	5	110.100000	279.742240	0.21790000	18853.10

CG311	CG331	HGA3	5	110.100000	279.742240	0.21790000	18853.10

CG321	CG331	HGA3	5	110.100000	289.532800	0.21790000	18853.10

NG2S0	CG331	HGA3	5	105.000000	418.400000	0.00000000	0.00

HGA3	CG331	HGA3	5	108.400000	297.064000	0.18020000	4518.72

CG2R53	CG3C51	CG3C52	5	106.500000	585.760000	0.00000000	0.00

CG2R53	CG3C51	SG311	5	103.000000	376.560000	0.00000000	0.00

CG2R53	CG3C51	HGA1	5	111.000000	485.344000	0.00000000	0.00

CG3C52	CG3C51	SG311	5	111.000000	920.480000	0.00000000	0.00

CG3C52	CG3C51	HGA1	5	111.400000	292.880000	0.21790000	18853.10

SG311	CG3C51	HGA1	5	111.300000	385.764800	0.00000000	0.00

CG2R53	CG3C52	CG3C51	5	106.500000	585.760000	0.00000000	0.00

CG2R53	CG3C52	HGA2	5	111.000000	485.344000	0.00000000	0.00

CG3C51	CG3C52	HGA2	5	111.400000	292.880000	0.21790000	18853.10

HGA2	CG3C52	HGA2	5	106.800000	322.168000	0.18020000	4518.72

CG2DC2	NG2P1	CG324	5	123.600000	560.656000	0.00000000	0.00

CG324	NG2P1	CG324	5	120.000000	521.326400	0.00000000	0.00

CG2R53	NG2R53	CG2R53	5	120.500000	460.240000	0.00000000	0.00

CG2R53	NG2R53	CG321	5	120.000000	418.400000	0.00000000	0.00

CG2O1	NG2S0	CG321	5	119.500000	351.456000	0.00000000	0.00

CG2O1	NG2S0	CG331	5	119.500000	351.456000	0.00000000	0.00

CG2O1	NG2S1	CG321	5	120.000000	418.400000	0.00000000	0.00

CG2O1	NG2S1	HGP1	5	123.000000	284.512000	0.00000000	0.00

CG321	NG2S1	HGP1	5	117.000000	292.880000	0.00000000	0.00

CG2R61	NG301	CG301	5	113.500000	401.664000	0.00000000	0.00

CG2R61	NG301	CG321	5	113.500000	401.664000	0.00000000	0.00

CG301	NG301	CG321	5	112.000000	585.760000	0.00000000	0.00

CG314	NG3P3	HGP2	5	109.500000	251.040000	0.20740000	16736.00

HGP2	NG3P3	HGP2	5	109.500000	368.192000	0.00000000	0.00

CG321	SG311	CG3C51	5	95.000000	284.512000	0.00000000	0.00

[ dihedraltypes ]

; i	j	k	l func	phi0	kphi mult

CG2DC2	CG2DC1	CG2DC1	CG301	9	180.000000	2.343040	1

CG2DC2	CG2DC1	CG2DC1	CG301	9	180.000000	29.288000	2

CG2DC2	CG2DC1	CG2DC1	HGA4	9	180.000000	21.756800	2

CG301	CG2DC1	CG2DC1	CG321	9	180.000000	41.840000	2

CG321	CG2DC1	CG2DC1	HGA4	9	180.000000	21.756800	2

CG2DC1	CG2DC1	CG2DC2	CG2DC1	9	180.000000	2.092000	1

CG2DC1	CG2DC1	CG2DC2	CG2DC1	9	0.000000	8.368000	2

CG2DC1	CG2DC1	CG2DC2	CG2DC1	9	0.000000	4.184000	3

CG2DC1	CG2DC1	CG2DC2	NG2P1	9	0.000000	2.092000	1

CG2DC1	CG2DC1	CG2DC2	NG2P1	9	180.000000	9.204800	2

CG2DC1	CG2DC1	CG2DC2	NG2P1	9	0.000000	4.602400	3

CG2DC1	CG2DC1	CG2DC2	NG2P1	9	0.000000	2.510400	4

CG301	CG2DC1	CG2DC2	CG2DC1	9	0.000000	3.765600	1

CG301	CG2DC1	CG2DC2	CG2DC1	9	180.000000	8.786400	2

CG301	CG2DC1	CG2DC2	CG2DC1	9	0.000000	0.920480	3

CG301	CG2DC1	CG2DC2	CG2DC1	9	180.000000	1.046000	5

CG301	CG2DC1	CG2DC2	CG2DC1	9	0.000000	0.418400	6

CG321	CG2DC1	CG2DC2	CG2DC1	9	180.000000	4.602400	1

CG321	CG2DC1	CG2DC2	CG2DC1	9	180.000000	2.928800	2

CG321	CG2DC1	CG2DC2	NG2P1	9	180.000000	4.602400	1

CG321	CG2DC1	CG2DC2	NG2P1	9	180.000000	2.928800	2

HGA4	CG2DC1	CG2DC2	CG2DC1	9	180.000000	4.184000	2

HGA4	CG2DC1	CG2DC2	CG2DC2	9	180.000000	4.184000	2

CG2DC1	CG2DC1	CG301	CG2R61	9	180.000000	0.000000	3

CG2DC2	CG2DC1	CG301	CG2R61	9	0.000000	0.000000	3

CG2DC2	CG2DC1	CG301	CG331	9	0.000000	1.255200	3

CG2DC1	CG2DC1	CG321	HGA2	9	0.000000	0.125520	3

CG2DC2	CG2DC1	CG321	CG321	9	0.000000	1.255200	3

CG2DC2	CG2DC1	CG321	HGA2	9	180.000000	1.255200	3

CG2DC1	CG2DC2	CG2DC2	CG2R61	9	180.000000	2.343040	1

CG2DC1	CG2DC2	CG2DC2	CG2R61	9	180.000000	29.288000	2

CG2DC2	CG2DC2	CG2R61	CG2R61	9	180.000000	3.138000	2

CG2DC2	CG2DC2	CG2R61	CG2R61	9	0.000000	0.794960	4

CG2R61	CG2DC2	CG2R61	CG2R61	9	180.000000	3.138000	2

CG2R61	CG2DC2	CG2R61	CG2R61	9	0.000000	0.794960	4

CG2DC1	CG2DC2	NG2P1	CG324	9	180.000000	29.288000	2

NG2S0	CG2O1	CG2R61	CG2R61	9	180.000000	4.184000	2

OG2D1	CG2O1	CG2R61	CG2R61	9	180.000000	4.184000	2

NG2S1	CG2O1	CG321	CG321	9	0.000000	0.000000	1

NG2S1	CG2O1	CG321	HGA2	9	0.000000	0.000000	3

OG2D1	CG2O1	CG321	CG321	9	180.000000	0.209200	6

OG2D1	CG2O1	CG321	HGA2	9	180.000000	0.000000	3

CG2R61	CG2O1	NG2S0	CG321	9	0.000000	6.694400	1

CG2R61	CG2O1	NG2S0	CG321	9	180.000000	16.736000	2

CG2R61	CG2O1	NG2S0	CG331	9	0.000000	6.694400	1

CG2R61	CG2O1	NG2S0	CG331	9	180.000000	16.736000	2

OG2D1	CG2O1	NG2S0	CG321	9	180.000000	10.878400	2

OG2D1	CG2O1	NG2S0	CG331	9	180.000000	10.878400	2

CG321	CG2O1	NG2S1	HGP1	9	180.000000	10.460000	2

OG2D1	CG2O1	NG2S1	CG321	9	180.000000	10.460000	2

OG2D1	CG2O1	NG2S1	HGP1	9	180.000000	10.460000	2

OG2D2	CG2O3	CG314	CG321	9	180.000000	0.209200	6

OG2D2	CG2O3	CG314	NG3P3	9	180.000000	13.388800	2

OG2D2	CG2O3	CG314	HGA1	9	180.000000	0.209200	6

NG2R53	CG2R53	CG3C51	CG3C52	9	180.000000	4.393200	3

NG2R53	CG2R53	CG3C51	SG311	9	180.000000	4.184000	3

NG2R53	CG2R53	CG3C51	HGA1	9	180.000000	0.000000	3

OG2D1	CG2R53	CG3C51	CG3C52	9	0.000000	0.334720	3

OG2D1	CG2R53	CG3C51	SG311	9	0.000000	0.334720	3

OG2D1	CG2R53	CG3C51	HGA1	9	0.000000	0.000000	3

NG2R53	CG2R53	CG3C52	CG3C51	9	180.000000	4.393200	3

NG2R53	CG2R53	CG3C52	HGA2	9	180.000000	0.000000	3

OG2D1	CG2R53	CG3C52	CG3C51	9	0.000000	0.334720	3

OG2D1	CG2R53	CG3C52	HGA2	9	0.000000	0.000000	3

CG3C51	CG2R53	NG2R53	CG2R53	9	180.000000	0.836800	2

CG3C51	CG2R53	NG2R53	CG321	9	180.000000	0.836800	2

CG3C52	CG2R53	NG2R53	CG2R53	9	180.000000	0.836800	2

CG3C52	CG2R53	NG2R53	CG321	9	180.000000	0.836800	2

OG2D1	CG2R53	NG2R53	CG2R53	9	180.000000	4.602400	2

OG2D1	CG2R53	NG2R53	CG321	9	180.000000	10.460000	2

CG2DC2	CG2R61	CG2R61	CG2O1	9	180.000000	12.970400	2

CG2DC2	CG2R61	CG2R61	CG2R61	9	180.000000	12.970400	2

CG2DC2	CG2R61	CG2R61	CG301	9	180.000000	12.970400	2

CG2DC2	CG2R61	CG2R61	HGR61	9	180.000000	10.041600	2

CG2O1	CG2R61	CG2R61	CG2R61	9	180.000000	12.970400	2

CG2O1	CG2R61	CG2R61	HGR61	9	180.000000	10.041600	2

CG2R61	CG2R61	CG2R61	CG2R61	9	180.000000	12.970400	2

CG2R61	CG2R61	CG2R61	CG301	9	180.000000	12.970400	2

CG2R61	CG2R61	CG2R61	CG311	9	180.000000	12.970400	2

CG2R61	CG2R61	CG2R61	NG301	9	180.000000	5.857600	2

CG2R61	CG2R61	CG2R61	HGR61	9	180.000000	17.572800	2

CG301	CG2R61	CG2R61	HGR61	9	180.000000	10.041600	2

CG311	CG2R61	CG2R61	NG301	9	180.000000	10.041600	2

CG311	CG2R61	CG2R61	HGR61	9	180.000000	10.041600	2

NG301	CG2R61	CG2R61	HGR61	9	180.000000	10.041600	2

HGR61	CG2R61	CG2R61	HGR61	9	180.000000	10.041600	2

CG2R61	CG2R61	CG301	CG2DC1	9	180.000000	0.962320	2

CG2R61	CG2R61	CG301	CG331	9	180.000000	0.962320	2

CG2R61	CG2R61	CG311	CG321	9	180.000000	0.962320	2

CG2R61	CG2R61	CG311	CG331	9	180.000000	0.962320	2

CG2R61	CG2R61	CG311	HGA1	9	180.000000	0.418400	6

CG2R61	CG2R61	NG301	CG301	9	180.000000	4.016640	2

CG2R61	CG2R61	NG301	CG301	9	0.000000	0.439320	4

CG2R61	CG2R61	NG301	CG301	9	180.000000	0.146440	6

CG2R61	CG2R61	NG301	CG321	9	180.000000	4.016640	2

CG2R61	CG2R61	NG301	CG321	9	0.000000	0.439320	4

CG2R61	CG2R61	NG301	CG321	9	180.000000	0.146440	6

CG331	CG301	CG321	CG311	9	0.000000	0.836800	3

CG331	CG301	CG321	HGA2	9	0.000000	0.815880	3

NG301	CG301	CG321	CG311	9	180.000000	0.669440	1

NG301	CG301	CG321	CG311	9	0.000000	1.631760	2

NG301	CG301	CG321	HGA2	9	0.000000	0.669440	3

CG2DC1	CG301	CG331	HGA3	9	0.000000	0.669440	3

CG2R61	CG301	CG331	HGA3	9	0.000000	0.167360	3

CG321	CG301	CG331	HGA3	9	0.000000	0.669440	3

CG331	CG301	CG331	HGA3	9	0.000000	0.669440	3

NG301	CG301	CG331	HGA3	9	0.000000	0.669440	3

CG321	CG301	NG301	CG2R61	9	180.000000	10.460000	1

CG321	CG301	NG301	CG2R61	9	0.000000	6.276000	2

CG321	CG301	NG301	CG2R61	9	0.000000	2.092000	3

CG321	CG301	NG301	CG321	9	180.000000	5.941280	1

CG321	CG301	NG301	CG321	9	0.000000	3.430880	2

CG321	CG301	NG301	CG321	9	0.000000	4.267680	3

CG331	CG301	NG301	CG2R61	9	180.000000	10.460000	1

CG331	CG301	NG301	CG2R61	9	0.000000	6.276000	2

CG331	CG301	NG301	CG2R61	9	0.000000	2.092000	3

CG331	CG301	NG301	CG321	9	180.000000	5.941280	1

CG331	CG301	NG301	CG321	9	0.000000	3.430880	2

CG331	CG301	NG301	CG321	9	0.000000	4.267680	3

CG2R61	CG311	CG321	CG301	9	0.000000	0.167360	3

CG2R61	CG311	CG321	HGA2	9	0.000000	0.000000	3

CG331	CG311	CG321	CG301	9	0.000000	0.836800	3

CG331	CG311	CG321	HGA2	9	0.000000	0.836800	3

HGA1	CG311	CG321	CG301	9	0.000000	0.815880	3

HGA1	CG311	CG321	HGA2	9	0.000000	0.815880	3

CG2R61	CG311	CG331	HGA3	9	0.000000	0.167360	3

CG321	CG311	CG331	HGA3	9	0.000000	0.836800	3

HGA1	CG311	CG331	HGA3	9	0.000000	0.815880	3

CG2O3	CG314	CG321	SG311	9	0.000000	0.836800	3

CG2O3	CG314	CG321	HGA2	9	0.000000	0.836800	3

NG3P3	CG314	CG321	SG311	9	0.000000	0.836800	3

NG3P3	CG314	CG321	HGA2	9	0.000000	0.836800	3

HGA1	CG314	CG321	SG311	9	0.000000	0.815880	3

HGA1	CG314	CG321	HGA2	9	0.000000	0.815880	3

CG2O3	CG314	NG3P3	HGP2	9	0.000000	0.418400	3

CG321	CG314	NG3P3	HGP2	9	0.000000	0.418400	3

HGA1	CG314	NG3P3	HGP2	9	0.000000	0.418400	3

CG2DC1	CG321	CG321	CG324	9	0.000000	0.794960	3

CG2DC1	CG321	CG321	HGA2	9	0.000000	0.794960	3

CG2O1	CG321	CG321	CG321	9	0.000000	0.815880	3

CG2O1	CG321	CG321	HGA2	9	0.000000	0.815880	3

CG321	CG321	CG321	NG2S0	9	0.000000	0.836800	3

CG321	CG321	CG321	HGA2	9	0.000000	0.815880	3

CG324	CG321	CG321	HGA2	9	0.000000	0.815880	3

NG2R53	CG321	CG321	NG2S1	9	0.000000	0.794960	3

NG2R53	CG321	CG321	HGA2	9	0.000000	0.815880	3

NG2S0	CG321	CG321	HGA2	9	0.000000	0.815880	3

NG2S1	CG321	CG321	HGA2	9	0.000000	0.815880	3

HGA2	CG321	CG321	HGA2	9	0.000000	0.920480	3

CG321	CG321	CG324	NG2P1	9	0.000000	0.815880	3

CG321	CG321	CG324	HGA2	9	0.000000	0.815880	3

HGA2	CG321	CG324	NG2P1	9	0.000000	0.815880	3

HGA2	CG321	CG324	HGA2	9	0.000000	0.815880	3

NG301	CG321	CG331	HGA3	9	0.000000	0.669440	3

HGA2	CG321	CG331	HGA3	9	0.000000	0.669440	3

CG321	CG321	NG2R53	CG2R53	9	0.000000	0.000000	1

HGA2	CG321	NG2R53	CG2R53	9	0.000000	0.000000	1

CG321	CG321	NG2S0	CG2O1	9	0.000000	7.531200	1

HGA2	CG321	NG2S0	CG2O1	9	0.000000	0.000000	3

CG321	CG321	NG2S1	CG2O1	9	0.000000	7.531200	1

CG321	CG321	NG2S1	HGP1	9	0.000000	0.000000	1

HGA2	CG321	NG2S1	CG2O1	9	0.000000	0.000000	3

HGA2	CG321	NG2S1	HGP1	9	0.000000	0.000000	3

CG331	CG321	NG301	CG2R61	9	180.000000	10.460000	1

CG331	CG321	NG301	CG2R61	9	0.000000	6.276000	2

CG331	CG321	NG301	CG2R61	9	0.000000	2.092000	3

CG331	CG321	NG301	CG301	9	180.000000	5.941280	1

CG331	CG321	NG301	CG301	9	0.000000	3.430880	2

CG331	CG321	NG301	CG301	9	0.000000	4.267680	3

HGA2	CG321	NG301	CG2R61	9	180.000000	0.000000	3

HGA2	CG321	NG301	CG301	9	0.000000	0.418400	3

CG314	CG321	SG311	CG3C51	9	0.000000	0.815880	3

HGA2	CG321	SG311	CG3C51	9	0.000000	1.171520	3

CG321	CG324	NG2P1	CG2DC2	9	0.000000	2.552240	1

CG321	CG324	NG2P1	CG2DC2	9	180.000000	2.594080	2

CG321	CG324	NG2P1	CG2DC2	9	0.000000	1.046000	3

CG321	CG324	NG2P1	CG2DC2	9	0.000000	2.510400	4

CG321	CG324	NG2P1	CG2DC2	9	0.000000	1.046000	6

CG321	CG324	NG2P1	CG324	9	180.000000	0.000000	6

HGA2	CG324	NG2P1	CG2DC2	9	180.000000	0.627600	3

HGA2	CG324	NG2P1	CG324	9	180.000000	0.000000	6

HGA3	CG331	NG2S0	CG2O1	9	0.000000	0.000000	3

HGA3	CG331	NG2S0	CG321	9	0.000000	1.757280	3

CG2R53	CG3C51	CG3C52	CG2R53	9	0.000000	0.000000	3

CG2R53	CG3C51	CG3C52	HGA2	9	0.000000	0.000000	3

SG311	CG3C51	CG3C52	CG2R53	9	180.000000	2.510400	3

SG311	CG3C51	CG3C52	HGA2	9	180.000000	0.627600	3

HGA1	CG3C51	CG3C52	CG2R53	9	0.000000	0.000000	3

HGA1	CG3C51	CG3C52	HGA2	9	0.000000	0.794960	3

HGA1	CG3C51	SG311	CG321	9	0.000000	1.171520	3

[ dihedraltypes ]

; 'improper' dihedrals

; i	j	k	l func	phi0	kphi

CG2DC2	CG2DC1	CG2DC1	NG2P1	2	0.000000 1004.160000

CG2O1	CG2R61	NG2S0	OG2D1	2	0.000000 1004.160000

CG2O1	CG321	NG2S1	OG2D1	2	0.000000 1004.160000

CG2O3	OG2D2	OG2D2	CG314	2	0.000000	803.328000

CG2R53	CG3C51	NG2R53	OG2D1	2	0.000000	753.120000

CG2R53	CG3C52	NG2R53	OG2D1	2	0.000000	753.120000

**PDB structure file**

ATOM	1 C1 ATT B 225	20.808	4.063 35.305 1.00 0.00	C

ATOM	2 C2 ATT B 225	22.116	3.969 35.007 1.00 0.00	C

ATOM	3 C3 ATT B 225	22.976	2.958 35.568 1.00 0.00	C

ATOM	4 N4 ATT B 225	24.263	3.011 35.572 1.00 0.00	N

ATOM	5 C5 ATT B 225	24.957	4.134 34.967 1.00 0.00	C

ATOM	6 C6 ATT B 225	24.195	4.771 33.795 1.00 0.00	C

ATOM	7 C7 ATT B 225	22.782	5.115 34.300 1.00 0.00	C

ATOM	8 C8 ATT B 225	25.151	1.876 35.817 1.00 0.00	C

ATOM	9 C9 ATT B 225	24.458	0.584 35.359 1.00 0.00	C

ATOM	10 C10 ATT B 225	23.077	0.556 36.015 1.00 0.00	C

ATOM	11 C11 ATT B 225	22.216	1.789 35.974 1.00 0.00	C

ATOM	12 C12 ATT B 225	20.920	1.870 36.335 1.00 0.00	C

ATOM	13 C13 ATT B 225	20.260	3.163 36.299 1.00 0.00	C

ATOM	14 C14 ATT B 225	19.025	3.388 36.774 1.00 0.00	C

ATOM	15 C15 ATT B 225	18.321	2.337 37.484 1.00 0.00	C

ATOM	16 C16 ATT B 225	18.713	1.033 37.338 1.00 0.00	C

ATOM	17 C17 ATT B 225	20.025	0.712 36.681 1.00 0.00	C

ATOM	18 C18 ATT B 225	19.663 -0.081 35.424 1.00 0.00	C

ATOM	19 C19 ATT B 225	20.672 -0.267 37.673 1.00 0.00	C

ATOM	20 C20 ATT B 225	17.817	0.095 37.808 1.00 0.00	C

ATOM	21 C21 ATT B 225	16.705	0.402 38.573 1.00 0.00	C

ATOM	22 C22 ATT B 225	16.512	1.729 38.892 1.00 0.00	C

ATOM	23 C23 ATT B 225	17.273	2.705 38.308 1.00 0.00	C

ATOM	24 C24 ATT B 225	15.425	2.230 39.795 1.00 0.00	C

ATOM	25 C25 ATT B 225	14.644	1.029 40.336 1.00 0.00	C

ATOM	26 C26 ATT B 225	14.584 -0.266 39.495 1.00 0.00	C

ATOM	27 N27 ATT B 225	15.975 -0.586 39.249 1.00 0.00	N

ATOM	28 C28 ATT B 225	16.319 -1.979 39.058 1.00 0.00	C

ATOM	29 C29 ATT B 225	16.041 -2.506 37.655 1.00 0.00	C

ATOM	30 C30 ATT B 225	13.719 -0.043 38.246 1.00 0.00	C

ATOM	31 C31 ATT B 225	13.930 -1.313 40.417 1.00 0.00	C

ATOM	32 C32 ATT B 225	14.516	3.122 38.929 1.00 0.00	C

ATOM	33 C33 ATT B 225	18.324	4.621 36.439 1.00 0.00	C

ATOM	34 C34 ATT B 225	17.453	4.716 35.380 1.00 0.00	C

ATOM	35 C35 ATT B 225	16.900	5.944 35.082 1.00 0.00	C

ATOM	36 C36 ATT B 225	17.138	7.026 35.886 1.00 0.00	C

ATOM	37 C37 ATT B 225	17.865	6.911 37.045 1.00 0.00	C

ATOM	38 C38 ATT B 225	18.522	5.709 37.238 1.00 0.00	C

ATOM	39 C39 ATT B 225	17.219	3.622 34.412 1.00 0.00	C

ATOM	40 O40 ATT B 225	18.141	3.193 33.725 1.00 0.00	O

ATOM	41 N41 ATT B 225	15.943	3.197 34.259 1.00 0.00	N

ATOM	42 C42 ATT B 225	14.846	3.578 35.100 1.00 0.00	C

ATOM	43 C43 ATT B 225	15.658	2.152 33.294 1.00 0.00	C

ATOM	44 C44 ATT B 225	15.528	0.849 34.084 1.00 0.00	C

ATOM	45 C45 ATT B 225	14.129	0.247 34.273 1.00 0.00	C

ATOM	46 C46 ATT B 225	13.484	0.094 32.930 1.00 0.00	C

ATOM	47 O47 ATT B 225	14.065 -0.214 31.885 1.00 0.00	O

ATOM	48 N48 ATT B 225	12.192	0.422 33.018 1.00 0.00	N

ATOM	49 C49 ATT B 225	11.223	0.226 31.985 1.00 0.00	C

ATOM	50 C50 ATT B 225	10.051	1.206 32.126 1.00 0.00	C

ATOM	51 N51 ATT B 225	9.123	1.108 31.046 1.00 0.00	N

ATOM	52 C52 ATT B 225	9.181	2.020 29.998 1.00 0.00	C

ATOM	53 O53 ATT B 225	9.731	3.117 30.067 1.00 0.00	O

ATOM	54 C54 ATT B 225	8.369	1.370 28.873 1.00 0.00	C

ATOM	55 C55 ATT B 225	7.673	0.204 29.543 1.00 0.00	C

ATOM	56 C56 ATT B 225	8.153	0.131 30.995 1.00 0.00	C

ATOM	57 O57 ATT B 225	8.054 -0.889 31.687 1.00 0.00	O

ATOM	58 S58 ATT B 225	8.269 -1.332 28.776 1.00 0.00	S

ATOM	59 C59 ATT B 225	7.096 -2.633 29.245 1.00 0.00	C

ATOM	60 CA ATT B 225	5.703 -2.448 28.625 1.00 0.00	C

ATOM	61 C	ATT B 225	4.531 -3.327 28.919 1.00 0.00	C

ATOM	62 O	ATT B 225	3.633 -3.448 28.082 1.00 0.00	O

ATOM	63 H63 ATT B 225	20.308	4.978 34.948 1.00 0.00	H

ATOM	64 H64 ATT B 225	25.854	3.739 34.477 1.00 0.00	H

ATOM	65 H65 ATT B 225	25.242	4.872 35.724 1.00 0.00	H

ATOM	66 H66 ATT B 225	24.731	5.664 33.413 1.00 0.00	H

ATOM	67 H67 ATT B 225	24.084	4.084 32.927 1.00 0.00	H

ATOM	68 H68 ATT B 225	22.165	5.512 33.472 1.00 0.00	H

ATOM	69 H69 ATT B 225	22.920	5.939 35.023 1.00 0.00	H

ATOM	70 H70 ATT B 225	26.094	1.961 35.259 1.00 0.00	H

ATOM	71 H71 ATT B 225	25.383	1.805 36.889 1.00 0.00	H

ATOM	72 H72 ATT B 225	25.043 -0.338 35.594 1.00 0.00	H

ATOM	73 H73 ATT B 225	24.360	0.731 34.264 1.00 0.00	H

ATOM	74 H74 ATT B 225	23.186	0.151 37.040 1.00 0.00	H

ATOM	75 H75 ATT B 225	22.489 -0.262 35.532 1.00 0.00	H

ATOM	76 H76 ATT B 225	19.207	0.538 34.617 1.00 0.00	H

ATOM	77 H77 ATT B 225	20.521 -0.547 34.896 1.00 0.00	H

ATOM	78 H78 ATT B 225	18.916 -0.884 35.599 1.00 0.00	H

ATOM	79 H79 ATT B 225	20.891	0.242 38.644 1.00 0.00	H

ATOM	80 H80 ATT B 225	21.534 -0.770 37.180 1.00 0.00	H

ATOM	81 H81 ATT B 225	20.004 -1.096 37.977 1.00 0.00	H

ATOM	82 H82 ATT B 225	18.005 -0.913 37.494 1.00 0.00	H

ATOM	83 H83 ATT B 225	17.034	3.760 38.369 1.00 0.00	H

ATOM	84 H84 ATT B 225	15.948	2.772 40.609 1.00 0.00	H

ATOM	85 H85 ATT B 225	13.595	1.330 40.511 1.00 0.00	H

ATOM	86 H86 ATT B 225	15.115	0.764 41.311 1.00 0.00	H

ATOM	87 H87 ATT B 225	17.403 -2.101 39.299 1.00 0.00	H

ATOM	88 H88 ATT B 225	15.874 -2.744 39.731 1.00 0.00	H

ATOM	89 H89 ATT B 225	15.921 -1.703 36.895 1.00 0.00	H

ATOM	90 H90 ATT B 225	16.905 -3.069 37.245 1.00 0.00	H

ATOM	91 H91 ATT B 225	15.185 -3.215 37.681 1.00 0.00	H

ATOM	92 H92 ATT B 225	12.687	0.267 38.528 1.00 0.00	H

ATOM	93 H93 ATT B 225	13.565 -1.080 37.882 1.00 0.00	H

ATOM	94 H94 ATT B 225	14.237	0.623 37.522 1.00 0.00	H

ATOM	95 H95 ATT B 225	12.904 -1.020 40.727 1.00 0.00	H

ATOM	96 H96 ATT B 225	14.534 -1.460 41.341 1.00 0.00	H

ATOM	97 H97 ATT B 225	13.935 -2.306 39.931 1.00 0.00	H

ATOM	98 H98 ATT B 225	13.479	3.049 39.334 1.00 0.00	H

ATOM	99 H99 ATT B 225	14.426	2.733 37.894 1.00 0.00	H

ATOM	100 0H10 ATT B 225	14.956	4.147 38.980 1.00 0.00	H

ATOM	101 1H10 ATT B 225	16.262	6.129 34.235 1.00 0.00	H

ATOM	102 2H10 ATT B 225	16.633	7.927 35.555 1.00 0.00	H

ATOM	103 3H10 ATT B 225	18.158	7.821 37.546 1.00 0.00	H

ATOM	104 4H10 ATT B 225	19.279	5.656 38.013 1.00 0.00	H

ATOM	105 5H10 ATT B 225	14.123	4.228 34.559 1.00 0.00	H

ATOM	106 6H10 ATT B 225	14.460	2.634 35.542 1.00 0.00	H

ATOM	107 7H10 ATT B 225	15.242	4.075 36.012 1.00 0.00	H

ATOM	108 8H10 ATT B 225	16.494	2.089 32.570 1.00 0.00	H

ATOM	109 9H10 ATT B 225	14.771	2.398 32.670 1.00 0.00	H

ATOM	110 0H11 ATT B 225	16.146	0.039 33.636 1.00 0.00	H

ATOM	111 1H11 ATT B 225	15.971	0.982 35.097 1.00 0.00	H

ATOM	112 2H11 ATT B 225	13.469	0.846 34.932 1.00 0.00	H

ATOM	113 3H11 ATT B 225	14.129 -0.767 34.723 1.00 0.00	H

ATOM	114 4H11 ATT B 225	11.887	0.766 33.904 1.00 0.00	H

ATOM	115 5H11 ATT B 225	11.804	0.605 31.116 1.00 0.00	H

ATOM	116 6H11 ATT B 225	10.802 -0.797 31.877 1.00 0.00	H

ATOM	117 7H11 ATT B 225	9.631	0.910 33.114 1.00 0.00	H

ATOM	118 8H11 ATT B 225	10.469	2.230 32.203 1.00 0.00	H

ATOM	119 9H11 ATT B 225	7.665	2.075 28.409 1.00 0.00	H

ATOM	120 0H12 ATT B 225	9.090	1.140 28.073 1.00 0.00	H

ATOM	121 1H12 ATT B 225	6.579	0.314 29.471 1.00 0.00	H

ATOM	122 2H12 ATT B 225	7.011 -2.780 30.345 1.00 0.00	H

ATOM	123 3H12 ATT B 225	7.535 -3.626 28.998 1.00 0.00	H

ATOM	124 HA ATT B 225	5.431 -1.496 29.063 1.00 0.00	H

ATOM	125 N	ATT B 225	5.857 -2.259 27.215 1.00 0.00	N

ATOM	126 HN ATT B 225	6.019 -3.103 26.713 1.00 0.00	H
