## Supplementary text 2. Topology for Alexa488-C5-maleimide. for "Sub-millisecond conformational dynamics of the A_2A_ adenosine receptor revealed by single-molecule FRET"

**GROMACS rtp file**

[ ALX ]

[ atoms ]

C1	CG2R61	-0.506	0

C2	CG2R61	0.065	2

C3	CG2R61	-0.110	3

C4	CG2R61	-0.109	4

C5	CG2R61	0.006	5

C6	CG2R61	0.410	6

O7	OG3R60	-0.410	7

C8	CG2R61	0.287	8

C9	CG2R61	-0.008	9

C10	CG2R67	-0.001	10

C11	CG2R67	0.010	11

C12	CG2R61	-0.066	12

C13	CG2R61	-0.115	13

C14	CG2R61	-0.064	14

C15	CG2R61	-0.101	15

C16	CG2R61	-0.101	16

C17	CG2O3	0.621	17

O18	OG2D2	-0.760	18

C19	CG2O1	0.433	19

O20	OG2D1	-0.428	20

N21	NG2S1	-0.566	21

C22	CG321	0.071	22

C23	CG321	-0.177	23

C24	CG321	-0.212	24

C25	CG321	-0.185	25

C26	CG321	0.048	26

N27	NG2R53	-0.167	27

C28	CG2R53	0.312	28

O29	OG2D1	-0.487	29

C30	CG3C52	-0.028	30

C31	CG3C51	0.160	31

C32	CG2R53	0.314	32

O33	OG2D1	-0.487	33

S34	SG311	-0.156	34

C35	CT2	-0.151	35

CA	CT1	0.383	36

C	C	0.448	37

O	O	-0.860	38

C39	CG2DC1	-0.140	39

C40	CG2DC1	-0.076	40

C41	CG2DC2	0.281	41

N42	NG2D1	-0.715	42

C43	CG2D1O	-0.603	43

S44	SG3O1	1.420	44

O45	OG2P1	-0.650	45

O46	OG2P1	-0.650	46

N47	NG2S3	-0.851	47

S48	SG3O1	1.340	48

O49	OG2P1	-0.650	49

O50	OG2P1	-0.650	50

H51	HGP1	0.326	51

H52	HGR61	0.115	52

H53	HGR61	0.115	53

H54	HGR61	0.115	54

H55	HGR61	0.115	55

H56	HGR61	0.115	56

H57	HGP1	0.314	57

H58	HGA2	0.090	58

H59	HGA2	0.090	59

H60	HGA2	0.090	60

H61	HGA2	0.090	61

H62	HGA2	0.090	62

H63	HGA2	0.090	63

H64	HGA2	0.090	64

H65	HGA2	0.090	65

H66	HGA2	0.090	66

H67	HGA2	0.090	67

H68	HGA2	0.090	68

H69	HGA2	0.090	69

H70	HGA1	0.090	70

H71	HA2	0.090	71

H72	HA2	0.090	72

HA	HB1	0.090	73

H76	HGA4	0.150	74

H77	HGA4	0.150	75

H78	HGP4	0.382	76

H79	HGP4	0.382	77

O80	OG2D2	-0.760	78

N	NH1	-0.358	79

HN	H	0.330	80

O84	OG2P1	-0.650	81

O85	OG2P1	-0.650	82

[ bonds ]

C1 C6

C1 C2

C1 S48

C2 C3

C2 N47

C3 C4

C3 H52

C4 C5

C4 H53

C5 C10

C5 C6

C6 O7

O7 C8

C8 C43

C8 C9

C9 C10

C9 C39

C10 C11

C11 C16

C11 C12

C12 C13

C12 H54

C13 C14

C13 C19

C14 C15

C14 H55

C15 C16

C15 H56

C16 C17

C17 O18

C17 O80

C19 O20

C19 N21

N21 C22

N21 H57

C22 C23

C22 H58

C22 H59

C23 C24

C23 H60

C23 H61

C24 C25

C24 H62

C24 H63

C25 C26

C25 H64

C25 H65

C26 N27

C26 H66

C26 H67

N27 C32

N27 C28

C28 O29

C28 C30

C30 C31

C30 H68

C30 H69

C31 C32

C31 S34

C31 H70

C32 O33

S34 C35

C35 CA

C35 H71

C35 H72

CA C

CA HA

CA N

C	O

C39 C40

C39 H76

C40 C41

C40 H77

C41 N42

C41 C43

N42 H51

C43 S44

S44 O45

S44 O46

S44 O84

N47 H78

N47 H79

S48 O49

S48 O50

S48 O85

N	HN

C +N

**GROMACS prm file**

[ bondtypes ]

; i	j func	b0	kb

CG2D1O	CG2DC2	1	0.14500000	251040.00

CG2D1O	CG2R61	1	0.14500000	305432.00

CG2D1O	SG3O1	1	0.17800000	192464.00

CG2DC1	CG2DC1	1	0.13400000	368192.00

CG2DC1	CG2DC2	1	0.14500000	251040.00

CG2DC1	CG2R61	1	0.14500000	305432.00

CG2DC1	HGA4	1	0.11000000	301666.40

CG2DC2	NG2D1	1	0.12760000	418400.00

CG2O1	CG2R61	1	0.14750000	251040.00

CG2O1	NG2S1	1	0.13450000	309616.00

CG2O1	OG2D1	1	0.12300000	518816.00

CG2O3	CG2R61	1	0.15000000	167360.00

CG2O3	CG314	1	0.15220000	167360.00

CG2O3	OG2D2	1	0.12600000	439320.00

CG2R53	CG3C51	1	0.15300000	251040.00

CG2R53	CG3C52	1	0.15300000	251040.00

CG2R53	NG2R53	1	0.13800000	384928.00

CG2R53	OG2D1	1	0.12350000	476976.00

CG2R61	CG2R61	1	0.13750000	255224.00

CG2R61	CG2R67	1	0.13750000	255224.00

CG2R61	NG2S3	1	0.13900000	334720.00

CG2R61	OG3R60	1	0.13500000	234304.00

CG2R61	SG3O1	1	0.17800000	192464.00

CG2R61	HGR61	1	0.10800000	284512.00

CG2R67	CG2R67	1	0.14900000	251040.00

CG314	CG321	1	0.15380000	186188.00

CG314	NG3P3	1	0.14800000	167360.00

CG314	HGA1	1	0.11110000	258571.20

CG321	CG321	1	0.15300000	186188.00

CG321	NG2R53	1	0.14300000	267776.00

CG321	NG2S1	1	0.14300000	267776.00

CG321	SG311	1	0.18180000	165686.40

CT2	SG311	1	0.18180000	165686.40 ;

CG321	HGA2	1	0.11110000	258571.20

CG3C51	CG3C52	1	0.15180000	163176.00

CG3C51	SG311	1	0.18180000	165686.40

CG3C51	HGA1	1	0.11000000	256897.60

CG3C52	HGA2	1	0.11000000	256897.60

NG2D1	HGP1	1	0.10000000	380744.00

NG2S1	HGP1	1	0.09970000	368192.00

NG2S3	HGP4	1	0.10000000	408358.40

NG3P3	HGP2	1	0.10400000	337230.40

OG2P1	SG3O1	1	0.14480000	451872.00

[ angletypes ]

; i	j	k func	theta0	ktheta	ub0	kub

CG2DC2	CG2D1O	CG2R61	5	117.200000	334.720000	0.00000000	0.00

CG2DC2	CG2D1O	SG3O1	5	113.000000	401.664000	0.00000000	0.00

CG2R61	CG2D1O	SG3O1	5	116.500000	167.360000	0.00000000	0.00

CG2DC1	CG2DC1	CG2DC2	5	123.000000	401.664000	0.00000000	0.00

CG2DC1	CG2DC1	CG2R61	5	122.000000	242.672000	0.00000000	0.00

CG2DC1	CG2DC1	HGA4	5	119.000000	351.456000	0.00000000	0.00

CG2DC2	CG2DC1	HGA4	5	118.000000	351.456000	0.00000000	0.00

CG2R61	CG2DC1	HGA4	5	120.000000	267.776000	0.00000000	0.00

CG2D1O	CG2DC2	CG2DC1	5	113.000000	401.664000	0.00000000	0.00

CG2D1O	CG2DC2	NG2D1	5	117.000000	468.608000	0.00000000	0.00

CG2DC1	CG2DC2	NG2D1	5	117.000000	468.608000	0.00000000	0.00

CG2R61	CG2O1	NG2S1	5	116.500000	669.440000	0.00000000	0.00

CG2R61	CG2O1	OG2D1	5	121.000000	251.040000	0.00000000	0.00

NG2S1	CG2O1	OG2D1	5	122.500000	669.440000	0.00000000	0.00

CG2R61	CG2O3	OG2D2	5	116.000000	334.720000	0.23530000	41840.00

CG314	CG2O3	OG2D2	5	116.000000	334.720000	0.23530000	41840.00

OG2D2	CG2O3	OG2D2	5	128.000000	836.800000	0.22587000	58576.00

CG3C51	CG2R53	NG2R53	5	105.500000 1004.160000	0.00000000	0.00

CG3C51	CG2R53	OG2D1	5	126.700000	543.920000	0.00000000	0.00

CG3C52	CG2R53	NG2R53	5	105.500000 1004.160000	0.00000000	0.00

CG3C52	CG2R53	OG2D1	5	126.700000	543.920000	0.00000000	0.00

NG2R53	CG2R53	OG2D1	5	127.800000	543.920000	0.00000000	0.00

CG2D1O	CG2R61	CG2R61	5	120.000000	301.248000	0.00000000	0.00

CG2D1O	CG2R61	OG3R60	5	113.900000	167.360000	0.00000000	0.00

CG2DC1	CG2R61	CG2R61	5	120.000000	301.248000	0.00000000	0.00

CG2DC1	CG2R61	CG2R67	5	120.000000	301.248000	0.00000000	0.00

CG2O1	CG2R61	CG2R61	5	119.000000	376.560000	0.00000000	0.00

CG2O3	CG2R61	CG2R61	5	119.000000	376.560000	0.00000000	0.00

CG2O3	CG2R61	CG2R67	5	119.000000	376.560000	0.00000000	0.00

CG2R61	CG2R61	CG2R61	5	120.000000	334.720000	0.24162000	29288.00

CG2R61	CG2R61	CG2R67	5	120.000000	334.720000	0.00000000	0.00

CG2R61	CG2R61	NG2S3	5	121.000000	502.080000	0.00000000	0.00

CG2R61	CG2R61	OG3R60	5	120.000000	334.720000	0.00000000	0.00

CG2R61	CG2R61	SG3O1	5	122.300000	83.680000	0.00000000	0.00

CG2R61	CG2R61	HGR61	5	120.000000	251.040000	0.21525000	18409.60

CG2R67	CG2R61	HGR61	5	120.000000	251.040000	0.00000000	0.00

CG2R61	CG2R67	CG2R61	5	120.000000	334.720000	0.00000000	0.00

CG2R61	CG2R67	CG2R67	5	120.000000	334.720000	0.00000000	0.00

CG2O3	CG314	CG321	5	108.000000	435.136000	0.00000000	0.00

CG2O3	CG314	NG3P3	5	110.000000	365.681600	0.00000000	0.00

CG2O3	CG314	HGA1	5	109.500000	418.400000	0.00000000	0.00

CG321	CG314	NG3P3	5	110.000000	566.513600	0.00000000	0.00

CG321	CG314	HGA1	5	110.100000	288.696000	0.21790000	18853.10

NG3P3	CG314	HGA1	5	107.500000	430.952000	0.00000000	0.00

CG314	CG321	SG311	5	112.500000	485.344000	0.00000000	0.00

CT1	CT2	SG311	5	112.500000	485.344000	0.00000000	0.00 ;

CG314	CG321	HGA2	5	110.100000	279.742240	0.21790000	18853.10

CG321	CG321	CG321	5	113.600000	488.272800	0.25610000	9338.69

CG321	CG321	NG2R53	5	113.500000	585.760000	0.00000000	0.00

CG321	CG321	NG2S1	5	113.500000	585.760000	0.00000000	0.00

CG321	CG321	HGA2	5	110.100000	221.752000	0.21790000	18853.10

NG2R53	CG321	HGA2	5	109.500000	430.952000	0.00000000	0.00

NG2S1	CG321	HGA2	5	109.500000	430.952000	0.00000000	0.00

SG311	CG321	HGA2	5	111.300000	385.764800	0.00000000	0.00

SG311	CT2	HA2	5	111.300000	385.764800	0.00000000	0.00 ;

HGA2	CG321	HGA2	5	109.000000	297.064000	0.18020000	4518.72

CG2R53	CG3C51	CG3C52	5	106.500000	585.760000	0.00000000	0.00

CG2R53	CG3C51	SG311	5	103.000000	376.560000	0.00000000	0.00

CG2R53	CG3C51	HGA1	5	111.000000	485.344000	0.00000000	0.00

CG3C52	CG3C51	SG311	5	111.000000	920.480000	0.00000000	0.00

CG3C52	CG3C51	HGA1	5	111.400000	292.880000	0.21790000	18853.10

SG311	CG3C51	HGA1	5	111.300000	385.764800	0.00000000	0.00

CG2R53	CG3C52	CG3C51	5	106.500000	585.760000	0.00000000	0.00

CG2R53	CG3C52	HGA2	5	111.000000	485.344000	0.00000000	0.00

CG3C51	CG3C52	HGA2	5	111.400000	292.880000	0.21790000	18853.10

HGA2	CG3C52	HGA2	5	106.800000	322.168000	0.18020000	4518.72

CG2DC2	NG2D1	HGP1	5	111.000000	623.416000	0.00000000	0.00

CG2R53	NG2R53	CG2R53	5	120.500000	460.240000	0.00000000	0.00

CG2R53	NG2R53	CG321	5	120.000000	418.400000	0.00000000	0.00

CG2O1	NG2S1	CG321	5	120.000000	418.400000	0.00000000	0.00

CG2O1	NG2S1	HGP1	5	123.000000	284.512000	0.00000000	0.00

CG321	NG2S1	HGP1	5	117.000000	292.880000	0.00000000	0.00

CG2R61	NG2S3	HGP4	5	111.600000	502.080000	0.00000000	0.00

HGP4	NG2S3	HGP4	5	117.000000	259.408000	0.00000000	0.00

CG314	NG3P3	HGP2	5	109.500000	251.040000	0.20740000	16736.00

HGP2	NG3P3	HGP2	5	109.500000	368.192000	0.00000000	0.00

CG2R61	OG3R60	CG2R61	5	115.000000	334.720000	0.00000000	0.00

CG321	SG311	CG3C51	5	95.000000	284.512000	0.00000000	0.00

CT2	SG311	CG3C51	5	95.000000	284.512000	0.00000000	0.00 ;

CG2D1O	SG3O1	OG2P1	5	98.000000	711.280000	0.00000000	0.00

CG2R61	SG3O1	OG2P1	5	98.000000	711.280000	0.00000000	0.00

OG2P1	SG3O1	OG2P1	5	109.470000 1087.840000	0.24500000	29288.00

[ dihedraltypes ]

; i	j	k	l func	phi0	kphi mult

CG2R61	CG2D1O	CG2DC2	CG2DC1	9	180.000000	2.092000	1

CG2R61	CG2D1O	CG2DC2	CG2DC1	9	0.000000	8.368000	2

CG2R61	CG2D1O	CG2DC2	CG2DC1	9	0.000000	4.184000	3

CG2R61	CG2D1O	CG2DC2	NG2D1	9	0.000000	2.092000	1

CG2R61	CG2D1O	CG2DC2	NG2D1	9	180.000000	9.204800	2

CG2R61	CG2D1O	CG2DC2	NG2D1	9	0.000000	4.602400	3

CG2R61	CG2D1O	CG2DC2	NG2D1	9	0.000000	2.510400	4

SG3O1	CG2D1O	CG2DC2	CG2DC1	9	180.000000	4.602400	1

SG3O1	CG2D1O	CG2DC2	CG2DC1	9	180.000000	2.928800	2

SG3O1	CG2D1O	CG2DC2	NG2D1	9	180.000000	4.602400	1

SG3O1	CG2D1O	CG2DC2	NG2D1	9	180.000000	2.928800	2

CG2DC2	CG2D1O	CG2R61	CG2R61	9	180.000000	3.138000	2

CG2DC2	CG2D1O	CG2R61	CG2R61	9	0.000000	0.794960	4

CG2DC2	CG2D1O	CG2R61	OG3R60	9	180.000000	3.138000	2

CG2DC2	CG2D1O	CG2R61	OG3R60	9	0.000000	0.794960	4

SG3O1	CG2D1O	CG2R61	CG2R61	9	180.000000	1.129680	2

SG3O1	CG2D1O	CG2R61	OG3R60	9	180.000000	1.129680	2

CG2DC2	CG2D1O	SG3O1	OG2P1	9	0.000000	0.016736	6

CG2R61	CG2D1O	SG3O1	OG2P1	9	0.000000	0.016736	6

CG2DC2	CG2DC1	CG2DC1	CG2R61	9	180.000000	2.343040	1

CG2DC2	CG2DC1	CG2DC1	CG2R61	9	180.000000	29.288000	2

CG2DC2	CG2DC1	CG2DC1	HGA4	9	180.000000	21.756800	2

CG2R61	CG2DC1	CG2DC1	HGA4	9	180.000000	21.756800	2

HGA4	CG2DC1	CG2DC1	HGA4	9	180.000000	21.756800	2

CG2DC1	CG2DC1	CG2DC2	CG2D1O	9	180.000000	6.276000	1

CG2DC1	CG2DC1	CG2DC2	CG2D1O	9	180.000000	4.184000	2

CG2DC1	CG2DC1	CG2DC2	CG2D1O	9	0.000000	6.276000	3

CG2DC1	CG2DC1	CG2DC2	NG2D1	9	0.000000	2.092000	1

CG2DC1	CG2DC1	CG2DC2	NG2D1	9	180.000000	9.204800	2

CG2DC1	CG2DC1	CG2DC2	NG2D1	9	0.000000	4.602400	3

CG2DC1	CG2DC1	CG2DC2	NG2D1	9	0.000000	2.510400	4

HGA4	CG2DC1	CG2DC2	CG2D1O	9	180.000000	4.184000	2

HGA4	CG2DC1	CG2DC2	NG2D1	9	180.000000	4.184000	2

CG2DC1	CG2DC1	CG2R61	CG2R61	9	180.000000	3.138000	2

CG2DC1	CG2DC1	CG2R61	CG2R61	9	0.000000	0.794960	4

CG2DC1	CG2DC1	CG2R61	CG2R67	9	180.000000	3.138000	2

CG2DC1	CG2DC1	CG2R61	CG2R67	9	0.000000	0.794960	4

HGA4	CG2DC1	CG2R61	CG2R61	9	180.000000	2.510400	2

HGA4	CG2DC1	CG2R61	CG2R67	9	180.000000	2.510400	2

CG2D1O	CG2DC2	NG2D1	HGP1	9	180.000000	50.208000	2

CG2DC1	CG2DC2	NG2D1	HGP1	9	180.000000	50.208000	2

NG2S1	CG2O1	CG2R61	CG2R61	9	180.000000	4.184000	2

OG2D1	CG2O1	CG2R61	CG2R61	9	180.000000	4.184000	2

CG2R61	CG2O1	NG2S1	HGP1	9	180.000000	10.460000	2

OG2D1	CG2O1	NG2S1	CG321	9	180.000000	10.460000	2

OG2D1	CG2O1	NG2S1	HGP1	9	180.000000	10.460000	2

OG2D2	CG2O3	CG2R61	CG2R61	9	180.000000	12.970400	2

OG2D2	CG2O3	CG2R61	CG2R67	9	180.000000	12.970400	2

OG2D2	CG2O3	CG314	CG321	9	180.000000	0.209200	6

OG2D2	CG2O3	CG314	NG3P3	9	180.000000	13.388800	2

OG2D2	CG2O3	CG314	HGA1	9	180.000000	0.209200	6

NG2R53	CG2R53	CG3C51	CG3C52	9	180.000000	4.393200	3

NG2R53	CG2R53	CG3C51	SG311	9	180.000000	4.184000	3

NG2R53	CG2R53	CG3C51	HGA1	9	180.000000	0.000000	3

OG2D1	CG2R53	CG3C51	CG3C52	9	0.000000	0.334720	3

OG2D1	CG2R53	CG3C51	SG311	9	0.000000	0.334720	3

OG2D1	CG2R53	CG3C51	HGA1	9	0.000000	0.000000	3

NG2R53	CG2R53	CG3C52	CG3C51	9	180.000000	4.393200	3

NG2R53	CG2R53	CG3C52	HGA2	9	180.000000	0.000000	3

OG2D1	CG2R53	CG3C52	CG3C51	9	0.000000	0.334720	3

OG2D1	CG2R53	CG3C52	HGA2	9	0.000000	0.000000	3

CG3C51	CG2R53	NG2R53	CG2R53	9	180.000000	0.836800	2

CG3C51	CG2R53	NG2R53	CG321	9	180.000000	0.836800	2

CG3C52	CG2R53	NG2R53	CG2R53	9	180.000000	0.836800	2

CG3C52	CG2R53	NG2R53	CG321	9	180.000000	0.836800	2

OG2D1	CG2R53	NG2R53	CG2R53	9	180.000000	4.602400	2

OG2D1	CG2R53	NG2R53	CG321	9	180.000000	10.460000	2

CG2D1O	CG2R61	CG2R61	CG2DC1	9	180.000000	12.970400	2

CG2D1O	CG2R61	CG2R61	CG2R67	9	180.000000	12.970400	2

CG2DC1	CG2R61	CG2R61	OG3R60	9	180.000000	12.970400	2

CG2O1	CG2R61	CG2R61	CG2R61	9	180.000000	12.970400	2

CG2O1	CG2R61	CG2R61	CG2R67	9	180.000000	12.970400	2

CG2O1	CG2R61	CG2R61	HGR61	9	180.000000	10.041600	2

CG2O3	CG2R61	CG2R61	CG2R61	9	180.000000	12.970400	2

CG2O3	CG2R61	CG2R61	HGR61	9	180.000000	10.041600	2

CG2R61	CG2R61	CG2R61	CG2R61	9	180.000000	12.970400	2

CG2R61	CG2R61	CG2R61	CG2R67	9	180.000000	12.970400	2

CG2R61	CG2R61	CG2R61	NG2S3	9	180.000000	20.920000	2

CG2R61	CG2R61	CG2R61	OG3R60	9	180.000000	12.970400	2

CG2R61	CG2R61	CG2R61	SG3O1	9	180.000000	12.970400	2

CG2R61	CG2R61	CG2R61	HGR61	9	180.000000	17.572800	2

CG2R67	CG2R61	CG2R61	OG3R60	9	180.000000	12.970400	2

CG2R67	CG2R61	CG2R61	HGR61	9	180.000000	17.572800	2

NG2S3	CG2R61	CG2R61	SG3O1	9	180.000000	12.970400	2

NG2S3	CG2R61	CG2R61	HGR61	9	180.000000	10.041600	2

OG3R60	CG2R61	CG2R61	SG3O1	9	180.000000	10.041600	2

HGR61	CG2R61	CG2R61	HGR61	9	180.000000	10.041600	2

CG2DC1	CG2R61	CG2R67	CG2R61	9	180.000000	12.970400	2

CG2DC1	CG2R61	CG2R67	CG2R67	9	180.000000	12.970400	2

CG2O3	CG2R61	CG2R67	CG2R61	9	180.000000	12.970400	2

CG2O3	CG2R61	CG2R67	CG2R67	9	180.000000	12.970400	2

CG2R61	CG2R61	CG2R67	CG2R61	9	180.000000	12.970400	2

CG2R61	CG2R61	CG2R67	CG2R67	9	180.000000	12.970400	2

HGR61	CG2R61	CG2R67	CG2R61	9	180.000000	17.572800	2

HGR61	CG2R61	CG2R67	CG2R67	9	180.000000	17.572800	2

CG2R61	CG2R61	NG2S3	HGP4	9	180.000000	5.648400	2

CG2D1O	CG2R61	OG3R60	CG2R61	9	0.000000	3.179840	2

CG2R61	CG2R61	OG3R60	CG2R61	9	0.000000	3.179840	2

CG2R61	CG2R61	SG3O1	OG2P1	9	0.000000	0.016736	6

CG2R61	CG2R67	CG2R67	CG2R61	9	180.000000	3.723760	2

CG2O3	CG314	CG321	SG311	9	0.000000	0.836800	3

CG2O3	CG314	CG321	HGA2	9	0.000000	0.836800	3

NG3P3	CG314	CG321	SG311	9	0.000000	0.836800	3

NG3P3	CG314	CG321	HGA2	9	0.000000	0.836800	3

HGA1	CG314	CG321	SG311	9	0.000000	0.815880	3

HGA1	CG314	CG321	HGA2	9	0.000000	0.815880	3

CG2O3	CG314	NG3P3	HGP2	9	0.000000	0.418400	3

CG321	CG314	NG3P3	HGP2	9	0.000000	0.418400	3

HGA1	CG314	NG3P3	HGP2	9	0.000000	0.418400	3

CG321	CG321	CG321	NG2R53	9	0.000000	0.836800	3

CG321	CG321	CG321	NG2S1	9	0.000000	0.836800	3

CG321	CG321	CG321	HGA2	9	0.000000	0.815880	3

NG2R53	CG321	CG321	HGA2	9	0.000000	0.815880	3

NG2S1	CG321	CG321	HGA2	9	0.000000	0.815880	3

HGA2	CG321	CG321	HGA2	9	0.000000	0.920480	3

CG321	CG321	NG2R53	CG2R53	9	0.000000	0.000000	1

HGA2	CG321	NG2R53	CG2R53	9	0.000000	0.000000	1

CG321	CG321	NG2S1	CG2O1	9	0.000000	7.531200	1

CG321	CG321	NG2S1	HGP1	9	0.000000	0.000000	1

HGA2	CG321	NG2S1	CG2O1	9	0.000000	0.000000	3

HGA2	CG321	NG2S1	HGP1	9	0.000000	0.000000	3

CG314	CG321	SG311	CG3C51	9	0.000000	0.815880	3

HGA2	CG321	SG311	CG3C51	9	0.000000	1.171520	3

CT1	CT2	SG311	CG3C51	9	0.000000	0.815880	3 ;

HA2	CT2	SG311	CG3C51	9	0.000000	1.171520	3 ;

CG2R53	CG3C51	CG3C52	CG2R53	9	0.000000	0.000000	3

CG2R53	CG3C51	CG3C52	HGA2	9	0.000000	0.000000	3

SG311	CG3C51	CG3C52	CG2R53	9	180.000000	2.510400	3

SG311	CG3C51	CG3C52	HGA2	9	180.000000	0.627600	3

HGA1	CG3C51	CG3C52	CG2R53	9	0.000000	0.000000	3

HGA1	CG3C51	CG3C52	HGA2	9	0.000000	0.794960	3

CG2R53	CG3C51	SG311	CG321	9	180.000000	1.004160	1

CG2R53	CG3C51	SG311	CG321	9	0.000000	1.548080	3

CG2R53	CG3C51	SG311	CT2	9	180.000000	1.004160	1 ;

CG2R53	CG3C51	SG311	CT2	9	0.000000	1.548080	3 ;

CG3C52	CG3C51	SG311	CT2	9	180.000000	1.004160	1 ;

CG3C52	CG3C51	SG311	CT2	9	0.000000	1.548080	3 ;

CG3C52	CG3C51	SG311	CG321	9	180.000000	1.004160	1

CG3C52	CG3C51	SG311	CG321	9	0.000000	1.548080	3

HGA1	CG3C51	SG311	CG321	9	0.000000	1.171520	3

HGA1	CG3C51	SG311	CT2	9	0.000000	1.171520	3 ;

[ dihedraltypes ]

; 'improper' dihedrals

; i	j	k	l func	phi0	kphi

CG2DC2	CG2D1O	CG2DC1	NG2D1	2	0.000000 1004.160000

CG2O1	CG2R61	NG2S1	OG2D1	2	0.000000 1004.160000

CG2O3	OG2D2	OG2D2	CG2R61	2	0.000000	803.328000

CG2O3	OG2D2	OG2D2	CG314	2	0.000000	803.328000

CG2R53	CG3C51	NG2R53	OG2D1	2	0.000000	753.120000

CG2R53	CG3C52	NG2R53	OG2D1	2	0.000000	753.120000

NG2S3	HGP4	HGP4	CG2R61	2	0.000000	-20.920000

**PDB structure file**

ATOM	1 C1 ALX B 310	-31.192 -19.880 25.930 1.00 0.00	C

ATOM	2 C2 ALX B 310	-31.928 -19.265 26.916 1.00 0.00	C

ATOM	3 C3 ALX B 310	-31.783 -17.909 27.138 1.00 0.00	C

ATOM	4 C4 ALX B 310	-31.008 -17.194 26.260 1.00 0.00	C

ATOM	5 C5 ALX B 310	-30.266 -17.819 25.284 1.00 0.00	C

ATOM	6 C6 ALX B 310	-30.327 -19.189 25.122 1.00 0.00	C

ATOM	7 O7 ALX B 310	-29.644 -19.764 24.100 1.00 0.00	O

ATOM	8 C8 ALX B 310	-28.774 -19.081 23.335 1.00 0.00	C

ATOM	9 C9 ALX B 310	-28.651 -17.722 23.511 1.00 0.00	C

ATOM	10 C10 ALX B 310	-29.474 -17.095 24.422 1.00 0.00	C

ATOM	11 C11 ALX B 310	-29.252 -15.627 24.555 1.00 0.00	C

ATOM	12 C12 ALX B 310	-29.843 -14.895 23.553 1.00 0.00	C

ATOM	13 C13 ALX B 310	-29.886 -13.522 23.641 1.00 0.00	C

ATOM	14 C14 ALX B 310	-29.305 -12.916 24.742 1.00 0.00	C

ATOM	15 C15 ALX B 310	-28.576 -13.658 25.647 1.00 0.00	C

ATOM	16 C16 ALX B 310	-28.518 -15.024 25.558 1.00 0.00	C

ATOM	17 C17 ALX B 310	-27.613 -15.720 26.528 1.00 0.00	C

ATOM	18 O18 ALX B 310	-26.792 -15.083 27.254 1.00 0.00	O

ATOM	19 C19 ALX B 310	-30.511 -12.742 22.560 1.00 0.00	C

ATOM	20 O20 ALX B 310	-31.143 -13.296 21.665 1.00 0.00	O

ATOM	21 N21 ALX B 310	-30.417 -11.400 22.563 1.00 0.00	N

ATOM	22 C22 ALX B 310	-31.156 -10.521 21.715 1.00 0.00	C

ATOM	23 C23 ALX B 310	-30.362 -9.211 21.605 1.00 0.00	C

ATOM	24 C24 ALX B 310	-30.860 -8.255 20.523 1.00 0.00	C

ATOM	25 C25 ALX B 310	-29.762 -7.319 20.025 1.00 0.00	C

ATOM	26 C26 ALX B 310	-29.411 -7.595 18.556 1.00 0.00	C

ATOM	27 N27 ALX B 310	-30.560 -7.600 17.705 1.00 0.00	N

ATOM	28 C28 ALX B 310	-31.834 -7.831 18.161 1.00 0.00	C

ATOM	29 O29 ALX B 310	-32.211 -7.852 19.339 1.00 0.00	O

ATOM	30 C30 ALX B 310	-32.766 -7.604 16.962 1.00 0.00	C

ATOM	31 C31 ALX B 310	-31.838 -7.177 15.834 1.00 0.00	C

ATOM	32 C32 ALX B 310	-30.416 -7.388 16.344 1.00 0.00	C

ATOM	33 O33 ALX B 310	-29.452 -6.853 15.795 1.00 0.00	O

ATOM	34 S34 ALX B 310	-32.048 -8.373 14.482 1.00 0.00	S

ATOM	35 C35 ALX B 310	-33.700 -8.128 13.758 1.00 0.00	C

ATOM	36 CA ALX B 310	-33.876 -8.356 12.248 1.00 0.00	C

ATOM	37 C	ALX B 310	-33.215 -7.291 11.430 1.00 0.00	C

ATOM	38 O	ALX B 310	-33.721 -6.219 11.114 1.00 0.00	O

ATOM	39 C39 ALX B 310	-27.637 -17.091 22.677 1.00 0.00	C

ATOM	40 C40 ALX B 310	-26.707 -17.806 22.034 1.00 0.00	C

ATOM	41 C41 ALX B 310	-26.692 -19.254 22.132 1.00 0.00	C

ATOM	42 N42 ALX B 310	-25.654 -19.952 21.846 1.00 0.00	N

ATOM	43 C43 ALX B 310	-27.959 -19.887 22.447 1.00 0.00	C

ATOM	44 S44 ALX B 310	-28.823 -20.869 21.241 1.00 0.00	S

ATOM	45 O45 ALX B 310	-28.471 -20.356 19.938 1.00 0.00	O

ATOM	46 O46 ALX B 310	-30.216 -20.546 21.487 1.00 0.00	O

ATOM	47 N47 ALX B 310	-32.807 -19.996 27.715 1.00 0.00	N

ATOM	48 S48 ALX B 310	-31.106 -21.654 25.836 1.00 0.00	S

ATOM	49 O49 ALX B 310	-29.777 -21.962 26.319 1.00 0.00	O

ATOM	50 O50 ALX B 310	-31.327 -21.926 24.432 1.00 0.00	O

ATOM	51 O80 ALX B 310	-27.612 -16.974 26.456 1.00 0.00	O

ATOM	52 N	ALX B 310	-33.678 -9.726 11.908 1.00 0.00	N

ATOM	53 O84 ALX B 310	-28.437 -22.218 21.567 1.00 0.00	O

ATOM	54 O85 ALX B 310	-32.185 -22.166 26.653 1.00 0.00	O

ATOM	55 H51 ALX B 310	-24.914 -19.308 21.650 1.00 0.00	H

ATOM	56 H52 ALX B 310	-32.349 -17.446 27.934 1.00 0.00	H

ATOM	57 H53 ALX B 310	-30.870 -16.125 26.382 1.00 0.00	H

ATOM	58 H54 ALX B 310	-30.406 -15.335 22.741 1.00 0.00	H

ATOM	59 H55 ALX B 310	-29.529 -11.862 24.845 1.00 0.00	H

ATOM	60 H56 ALX B 310	-28.103 -13.079 26.427 1.00 0.00	H

ATOM	61 H57 ALX B 310	-29.773 -11.068 23.254 1.00 0.00	H

ATOM	62 H58 ALX B 310	-32.245 -10.378 21.873 1.00 0.00	H

ATOM	63 H59 ALX B 310	-31.127 -11.026 20.728 1.00 0.00	H

ATOM	64 H60 ALX B 310	-29.374 -9.601 21.287 1.00 0.00	H

ATOM	65 H61 ALX B 310	-30.272 -8.801 22.636 1.00 0.00	H

ATOM	66 H62 ALX B 310	-31.746 -7.665 20.848 1.00 0.00	H

ATOM	67 H63 ALX B 310	-31.270 -8.964 19.758 1.00 0.00	H

ATOM	68 H64 ALX B 310	-30.261 -6.350 20.268 1.00 0.00	H

ATOM	69 H65 ALX B 310	-28.857 -7.247 20.668 1.00 0.00	H

ATOM	70 H66 ALX B 310	-28.737 -6.770 18.232 1.00 0.00	H

ATOM	71 H67 ALX B 310	-28.783 -8.502 18.461 1.00 0.00	H

ATOM	72 H68 ALX B 310	-33.370 -8.503 16.829 1.00 0.00	H

ATOM	73 H69 ALX B 310	-33.453 -6.762 17.153 1.00 0.00	H

ATOM	74 H70 ALX B 310	-32.101 -6.142 15.558 1.00 0.00	H

ATOM	75 H71 ALX B 310	-34.440 -8.632 14.417 1.00 0.00	H

ATOM	76 H72 ALX B 310	-34.005 -7.068 13.886 1.00 0.00	H

ATOM	77 HA ALX B 310	-34.897 -8.149 11.972 1.00 0.00	H

ATOM	78 H76 ALX B 310	-27.555 -15.991 22.591 1.00 0.00	H

ATOM	79 H77 ALX B 310	-25.906 -17.277 21.500 1.00 0.00	H

ATOM	80 H78 ALX B 310	-32.861 -20.951 27.447 1.00 0.00	H

ATOM	81 H79 ALX B 310	-32.682 -19.706 28.659 1.00 0.00	H

ATOM	82 HN ALX B 310	-33.381 -10.337 12.641 1.00 0.00	H
